# Combination of engineered cell type-specific promoters and a high-efficiency AAV capsid restores hearing in adult DFNB1 mice model with demonstrated safety in nonhuman primate

**DOI:** 10.64898/2026.04.15.718827

**Authors:** Shao Wei Hu, Cheng Ye, Guannan Geng, Yicheng Zeng, Yihan Bao, Sen Zhang, Chong Cui, Yingting Zhang, Dan Mu, Daqi Wang, Xintai Fan, Ziting Chen, Biyun Zhu, Shuang Han, Hongxing Wang, Qing Su, Lei Han, Xiaoting Hu, Honghai Tang, Xiangnan Wang, Zhuoer Sun, Haoyun Yu, Haotian Deng, Zimu Cai, Huawei Li, Hongbo Yang, Guangbin Sun, Yilai Shu

**Author notes:** Corresponding author. (Yilai Shu); (Guangbin Sun). These authors contributed equally to this work (Shao Wei Hu, Cheng Ye, Guannan Geng, Yicheng Zeng, Yihan Bao, Sen Zhang, Chong Cui).

## Abstract

A major challenge in gene therapy for *GJB2*-related hearing loss (DFNB1)—the most common form of hereditary deafness—is achieving efficient and precise connexin 26 delivery. Herein, we engineered two cell type-specific promoters (GJB2-1 and WFS1-2274) and developed an AAV capsid, AAV-MAS012, with enhanced transduction efficiency in mature cochlear cells. Our AAV-mediated gene therapy systems restored hearing of low-to-mid-frequencies in newborn *Gjb2* cKO mice to wild-type levels and maintained for 45 weeks. Additionally, our therapeutic systems restored low-to-mid-frequencies hearing function to wild-type levels in adult *Gjb2* cKO mice. A humanized version of the therapy, AAV-MAS012-WFS1-2274-hGJB2, rescued hearing function in two distinct *Gjb2*-deficient mouse models, and demonstrated a favorable safety profile in nonhuman primates. This study represents the first successful hearing restoration in adult *Gjb2*-deficient mice. The significant therapeutic efficacy of the humanized gene therapy system shows great potential for clinical translation in DFNB1 patients.

---

Hearing loss is a widespread public health problem and is the most prevalent sensory deficit disorder in humans^1–3^. The number of people suffering from disabling hearing impairment is about 466 million globally, accounting for 5% of the world’s total population^4–8^. To date, more than 8,000 mutation loci in about 200 deafness-causing genes have been identified^9, 10^. Pathogenic variants in *GJB2* (gap junction protein β2), the most common cause of autosomal recessive deafness 1 (DFNB1) in humans^11–13^, result in hearing loss ranging from mild to profound, show congenital or delayed onset, and can be either progressive or non-progressive^14, 15^. Approximately one-quarter of non-syndromic hereditary hearing loss cases and over 50% of autosomal recessive non-syndromic hereditary hearing loss cases are caused by pathogenic mutations in *GJB2*^12, 16^, but effective biological therapies remain unavailable. For individuals with hearing loss resulting from *GJB2* mutations, the primary treatment approach is cochlear implants. However, cochlear implants present several limitations, including the inability to fully restore normal hearing function, difficulties in processing complex auditory environments, and poor discrimination of background noise, and for many recipients, the visible external components of a cochlear implant contribute to experiences of shame and stigma^17–19^.

Gene therapy aims to transfer therapeutic transgenes into particular cells or organs in order to treat diseases at the genetic level^20^. Adeno-associated virus (AAV)-mediated gene therapy has successfully rescued hearing function and speech perception in autosomal recessive deafness 9 (DFNB9) patients^4, 5, 21–24^, offering the possibility and hope for gene therapy in the field of hereditary deafness. In the cochlea, the GJB2 gene encodes Connexin 26 (Cx26), which is mainly expressed in various supporting cells (SCs)—including pillar cells (PCs), Deiter’s cells (DCs), and Claudius cells (CCs)—within the cochlear epithelium, as well as in the stria vascularis (SV), the spiral ligament (SPL), and the spiral limbus (SL), yet is not expressed in auditory sensory hair cells (HCs)^25^. Over 340 pathogenic variants in *GJB2,* like c.235delC, c.35delG and c.167delT, have been described^26, 27^, and gene replacement thus might be an optimal strategy for treating DFNB1. However, gene replacement strategies for *GJB2* gene therapy face two major challenges. First, ectopic expression of Cx26 in the inner ear may induce inner hair cell (IHC) apoptosis and loss of hearing function^28, 29^. Second, there is a lack of an effective delivery system for adult inner ear *GJB2*-expressing cells^28–31^. In the field of gene therapy for *GJB2*-related genetic deafness, several therapeutic gene replacement systems incorporating cell type-specific promoters or enhancers have been constructed^28–30^. However, the therapeutic efficacy of these systems remains to be improved, particularly for achieving long-term hearing restoration and auditory recovery in adult *Gjb2*-deficient mouse models. Therefore, new cell type-specific promoters and new AAV serotypes that have greater transduction efficiency in the adult mammalian inner ear are urgently needed.

In this study, we employed multiple promoter construction strategies and successfully constructed two cell type-specific promoters—GJB2-1 and WFS1-2274—and validated their specificity and transcriptional efficiency. We also engineered a novel AAV capsid, AAV-MAS012, which exhibited high transduction efficiency in mature mammalian inner ear SCs. Delivered by AAV-ie, our *GJB2* gene replacement systems driven by these cell type-specific promoters successfully restored Cx26 expression in *Gjb2* conditional knockout (cKO) mice, prevented the degeneration of HCs and SCs, and led to the sustained recovery of hearing in neonatal *Gjb2* cKO mice for up to 45 weeks. Moreover, delivery of the human *GJB2* coding sequence driven by the WFS1-2274 promoter using AAV-MAS012 also restored hearing in adult *Gjb2* cKO mice and in *Gjb6*-null mutant mice. The development of these cell type-specific promoters, the high-efficiency AAV capsid, and the corresponding humanized *GJB2* gene replacement system represents a significant step toward the clinical translation of gene therapy for *GJB2*-related hereditary deafness.

## Results

### Toxicity of the *GJB2* gene therapy system using a ubiquitous promoter

It has been demonstrated that ectopic expression of Cx26 in the inner ear leads to HC apoptosis and hearing loss^12, 28, 29^. In our study, the administration of CAG-driven mouse *Gjb2* cDNA via AAV-ie capsids^32–34^ (AAV-ie-CAG-mGJB2) triggered complete hearing loss in adult wild-type (WT) mice within four weeks (Extended Data Fig. 1a), along with the total absence of IHCs (Extended Data Fig. 1b). In contrast, mice from the control group (AAV-ie-CAG-EGFP) exhibited normal hearing function (Extended Data Fig. 1a) and normal inner ear morphology (Extended Data Fig. 1c). These results align with prior findings showing that ectopic expression of Cx26 in the cochlea leads to apoptosis of IHCs and subsequent hearing loss^12, 29^, thus suggesting the need to engineer a cell type-specific promoter that can precisely restrict the expression of Cx26 protein in gene replacement strategy for *GJB2-*related hearing loss.

### Strategies for designing cell type-specific promoters and identification of the GJB2-1 and WSF1-2274 promoters

We created a library of 21 AAV plasmids, each equipped with a synthetic promoter sequence with different lengths positioned 5′ of a fluorescent marker. The promoter sequences were constructed using four different strategies (Fig. 1a). 1) We predicted a 3628 bp GJB2-1 and a 3600 bp GJB2-2 promoter sequence upstream of the start codon of the mouse *Gjb2* gene (Fig. 1a-1). 2) Based on RNA-seq data, a 2001 bp Y1 promoter, a 2901 bp Y2 promoter, and a 3312 bp Y3 promoter were constructed. A 380 bp CMV enhancer was added to Y1 and Y3 to yield a 2381 bp Y4 promoter and a 3592 bp Y5 promoter, respectively (Fig. 1a-2). 3) According to previous reports, we selected two human-origin GJB2 promoter sequences and modified them to create a hGJB2-2 promoter and a hGJB2-10 promoter^35, 36^ (Fig. 1a-3). 4) We selected six genes whose expression patterns are relatively consistent with that of *Gjb2* in the mouse inner ear (Supplementary Table 1)^37^, for each gene, we predicted a ≈ 3450 bp promoter, and a ≈ 2000 bp promoter^37–40^ (Fig. 1a-4). We then constructed different AAV plasmids with these 21 different promoters and packaged these recombinant plasmids into AAV-ie to drive the expression of fluorescent marker proteins.

**Fig. 1.**
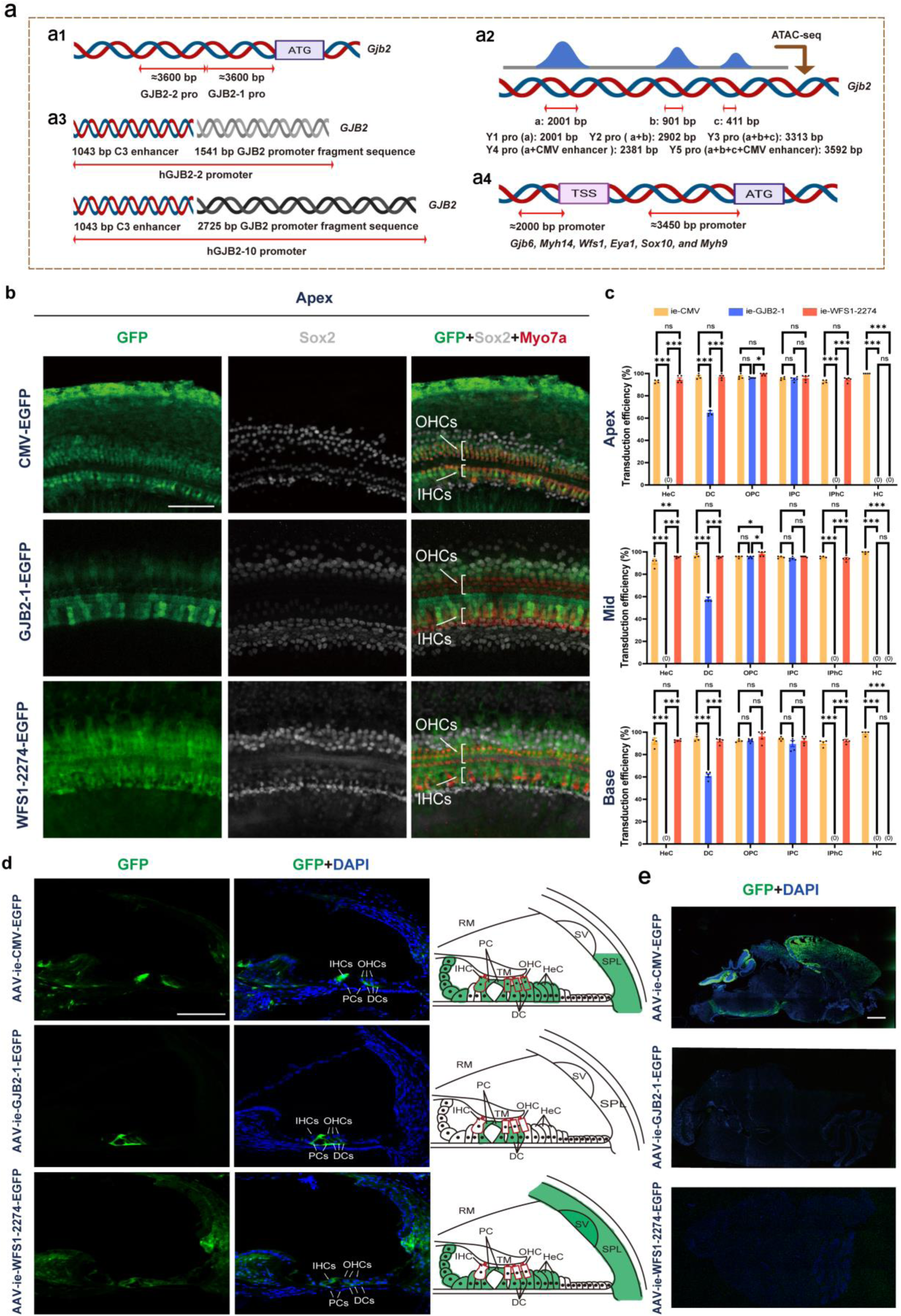
Generation of cochlear cell type-specific promoters. **a,** The different promoter design strategies used in this study. (**a-1**) We predicted a 3628 bp (–3628 bp to –1 bp) GJB2-1 and a 3600 bp (–7200 bp to –3601 bp) GJB2-2 promoter sequence upstream of the start codon of the mouse *Gjb2* gene. (**a-2**) Based on RNA-seq data, three sequences, including a 2001 bp sequence (sequence a), a 900 bp sequence (sequence b), and a 411 bp sequence (sequence c) within the mouse *Gjb2* gene have been predicted to be promoter sequences, and a 2001 bp Y1 promoter (sequence a), a 2901 bp Y2 promoter (sequence a+b), and a 3312 bp Y3 promoter (sequence a+b+c) were constructed. Of these three promoters, a 380 bp CMV enhancer was added to Y1 and Y3 to yield a 2381 bp Y4 promoter and a 3592 bp Y5 promoter, respectively. (**a-3**) According to previous reports, we identified several candidate human-origin GJB2 promoter sequences, and we selected two promoter sequences and modified them to create a hGJB2-2 promoter (a 1043 bp C3 enhancer + a 1541 bp human GJB2 promoter fragment sequence) and a hGJB2-10 promoter (1043 bp C3 enhancer + 2725 bp −2.725*hCx26*/CAT human *GJB2* promoter fragment sequence). (**a-4**) Based on the gene expression profiles and the localization of the encoded proteins associated with hereditary hearing loss, we selected six genes (*Gjb6*, *Myh14*, *Wfs1*, *Eya1*, *Sox10*, and *Myh9*) whose expression patterns are relatively consistent with that of *Gjb2* in the mouse inner ear. For each gene, we predicted two promoter regions: one spanning approximately 3450 bp upstream of the start codon (≈ –3450 bp to –1 bp), and another spanning approximately 2000 bp upstream of the transcription start site (≈ –2000 bp to –1 bp). **b,** The expression of GFP in the cochleae of AAV-ie-CMV-EGFP, AAV-ie-GJB2-1-EGFP, and AAV-ie-WFS1-2274-EGFP injected mice. **c,** The expression efficiency of GFP in HeCs, DCs, OPCs, IPCs, IPhCs, and HCs in the ears injected with AAV-ie-CMV-EGFP (n = 4), AAV-ie-GJB2-1-EGFP (n = 5), or AAV-ie-WFS1-2274-EGFP (n = 5). Scale bars: 100 μm. **d,** Immunofluorescent staining of cochlear cryosections (Left) and their schematic diagrams (Right) in the AAV-ie-CMV-EGFP, AAV-ie-GJB2-1-EGFP, and AAV-ie-WFS1-2274-EGFP groups. Scale bars: 100 μm. **e,** The expression of GFP in the central brain in the AAV-ie-CMV-EGFP, AAV-ie-GJB2-1-EGFP, and AAV-ie-WFS1-2274-EGFP groups. Scale bars: 1mm. Green: GFP; Gray: Sox2; Red: Myo7a; Blue: DAPI. ATG: translation start site, pro: promoter, ATAC-seq: assay for transposase-accessible chromatin sequencing, TSS: transcription start site.

**Fig. 2.**
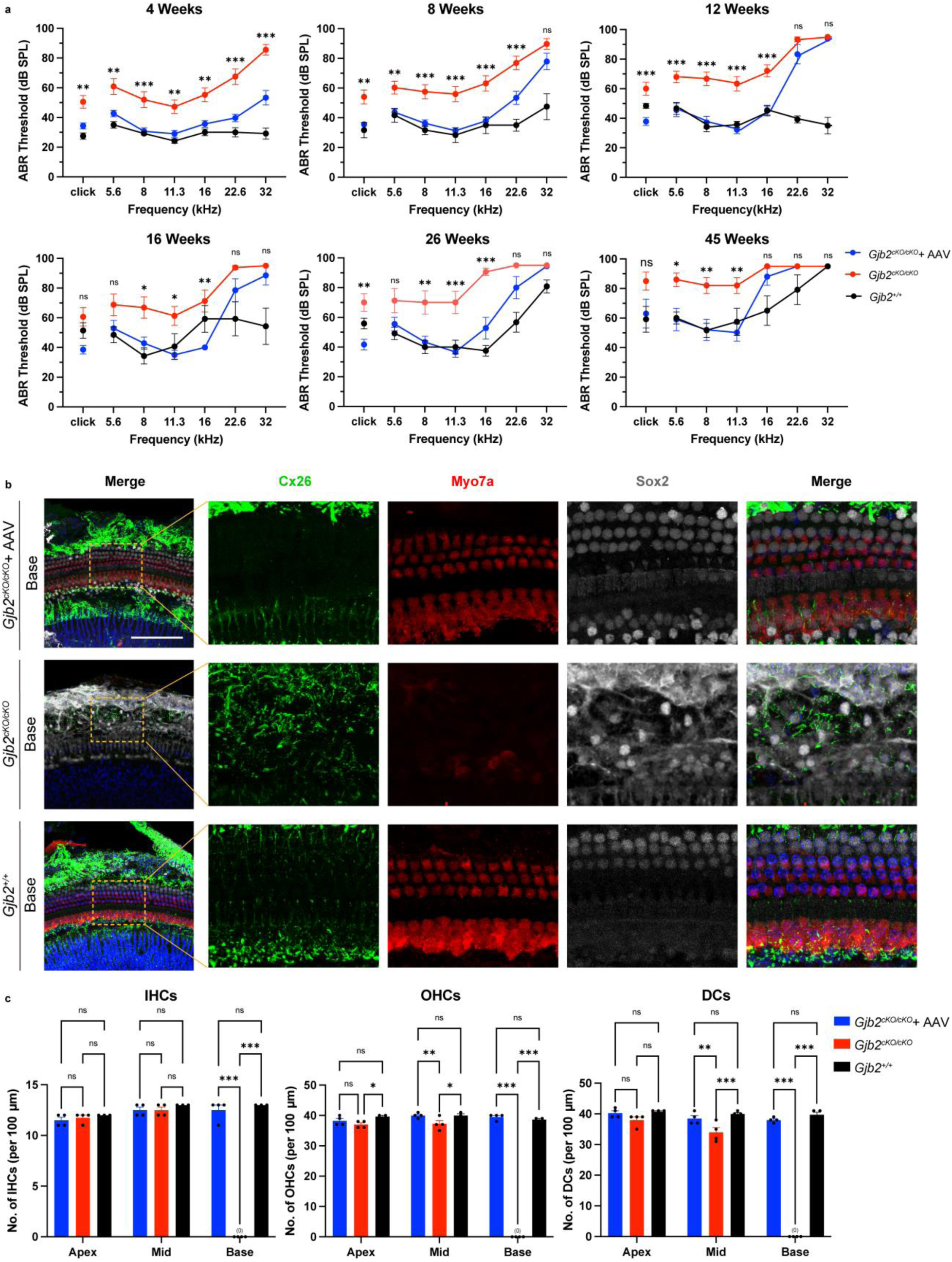
The expression of mouse Cx26 protein in the cochlear under the control of the GJB2-1 promoter restored hearing in newborn Fgfr3-iCreERT2; Cx26^loxP/loxP^ mice. **a,** The ABR thresholds of Fgfr3-iCreERT2; Cx26^loxP/loxP^ mice for click sound stimuli and pure-tone stimuli were recorded at 4 weeks (WT, n = 6; treated Fgfr3-iCreERT2; Cx26^loxP/loxP^ mice, n = 15, untreated Fgfr3-iCreERT2; Cx26^loxP/loxP^ mice, n = 18), 8 weeks (WT, n = 6; treated Fgfr3-iCreERT2; Cx26^loxP/loxP^ mice, n = 19, untreated Fgfr3-iCreERT2; Cx26^loxP/loxP^ mice, n = 16), 12 weeks (WT, n = 6; treated Fgfr3-iCreERT2; Cx26^loxP/loxP^ mice, n = 9, untreated Fgfr3-iCreERT2; Cx26^loxP/loxP^ mice, n = 12), 16 weeks (WT, n = 6; treated Fgfr3-iCreERT2; Cx26^loxP/loxP^ mice, n = 7, untreated Fgfr3-iCreERT2; Cx26^loxP/loxP^ mice, n = 8), 26 weeks (WT, n = 6; treated Fgfr3-iCreERT2; Cx26^loxP/loxP^ mice, n = 9, untreated Fgfr3-iCreERT2; Cx26^loxP/loxP^ mice, n = 8), and 45 weeks (WT, n = 6; treated Fgfr3-iCreERT2; Cx26^loxP/loxP^ mice, n = 5, untreated Fgfr3-iCreERT2; Cx26^loxP/loxP^ mice, n = 5) after therapeutic injection at P0–P2. **b,** The expression of Cx26 was decreased in PCs and DCs in the base turn of Fgfr3-iCreERT2; Cx26^loxP/loxP^ mice, and AAV-MAS012-GJB2-1-mGJB2 (titer: 2E13 VG/mL) injection restored the expression of mouse Cx26 in PCs and DCs in the base turn of Fgfr3-iCreERT2; Cx26^loxP/loxP^ mice. The dashed boxes highlight the regions with restored expression of mouse Cx26 protein, and the right-hand panels show zoomed-in views of the areas within the dashed boxes. **c,** AAV-MAS012-GJB2-1-mGJB2 injection inhibited cell death in IHCs, OHCs, and DCs in Fgfr3-iCreERT2; Cx26^loxP/loxP^ mice (WT, n = 3; treated Fgfr3-iCreERT2; Cx26^loxP/loxP^, n = 4; untreated Fgfr3-iCreERT2; Cx26^loxP/loxP^, n = 4). Green: Cx26; Red: Myo7a; Gray: Sox2; Blue: DAPI. Scale bars: 100 μm.

**Fig. 3.**
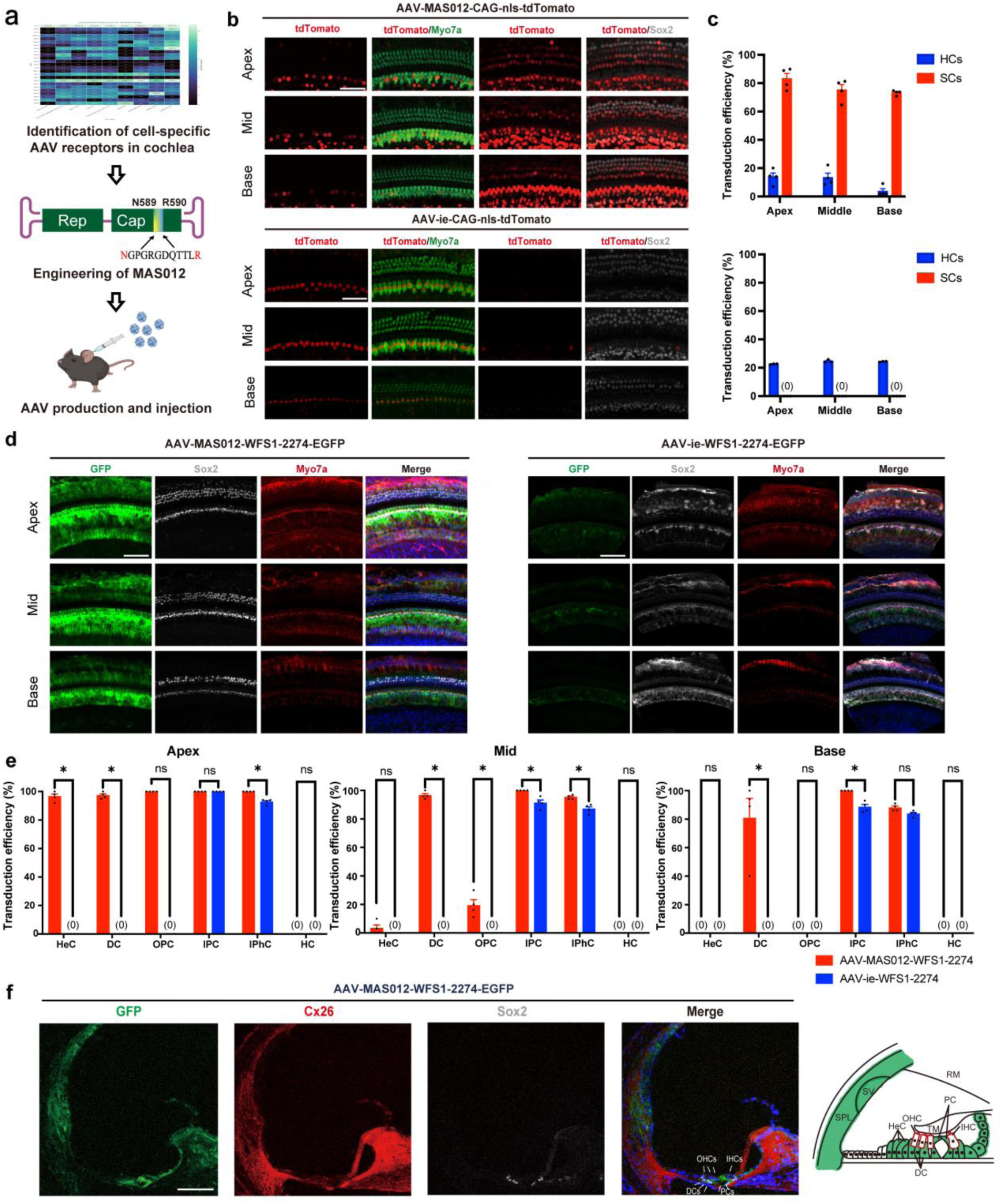
Engineering of a novel AAV capsid with high transduction efficiency in the mature inner ear. **a,** Workflow for identification of cell-specific receptors in cochlea and engineering of novel AAV capsid. **b,** The expression region of tdTomato in the injected ears of adult WT mice in the AAV-MAS012-CAG-tdTomato (titer: 1E13 VG/mL) and AAV-ie-CAG-tdTomato groups (titer: 1E13 VG/mL). Red: tdTomato; Green: Myo7a; Gray: Sox2. **c,** The expression efficiency of tdTomato in SCs and HCs of the injected ear in the AAV-MAS012-CAG-tdTomato (n = 4) and AAV-ie-CAG-tdTomato (n = 3) groups. **d,** The expression region of GFP in the injected ears of adult WT mice in the AAV-MAS012-WFS1-2274-EGFP (titer: 1E13 VG/mL) and AAV-ie-WFS1-2274-EGFP (titer: 1E13 VG/mL) groups, respectively. Green: GFP; Gray: Sox2; Red: Myo7a; Blue: DAPI. **e,** The expression efficiency of GFP in HeCs, DCs, OPCs, IPCs, IPhCs, and HCs of the injected ear in the AAV-MAS012-WFS1-2274-EGFP (n = 4) and AAV-ie-WFS1-2274-EGFP (n = 4) groups. **f,** Immunofluorescent staining of cochlear cryosections in AAV-MAS012-WFS1-2274-EGFP–treated adult WT mice (Left) and their schematic diagrams (Right). Green: GFP; Gray: Sox2; Red: Cx26; Blue: DAPI. Scale bars: 100 μm.

**Fig. 4.**
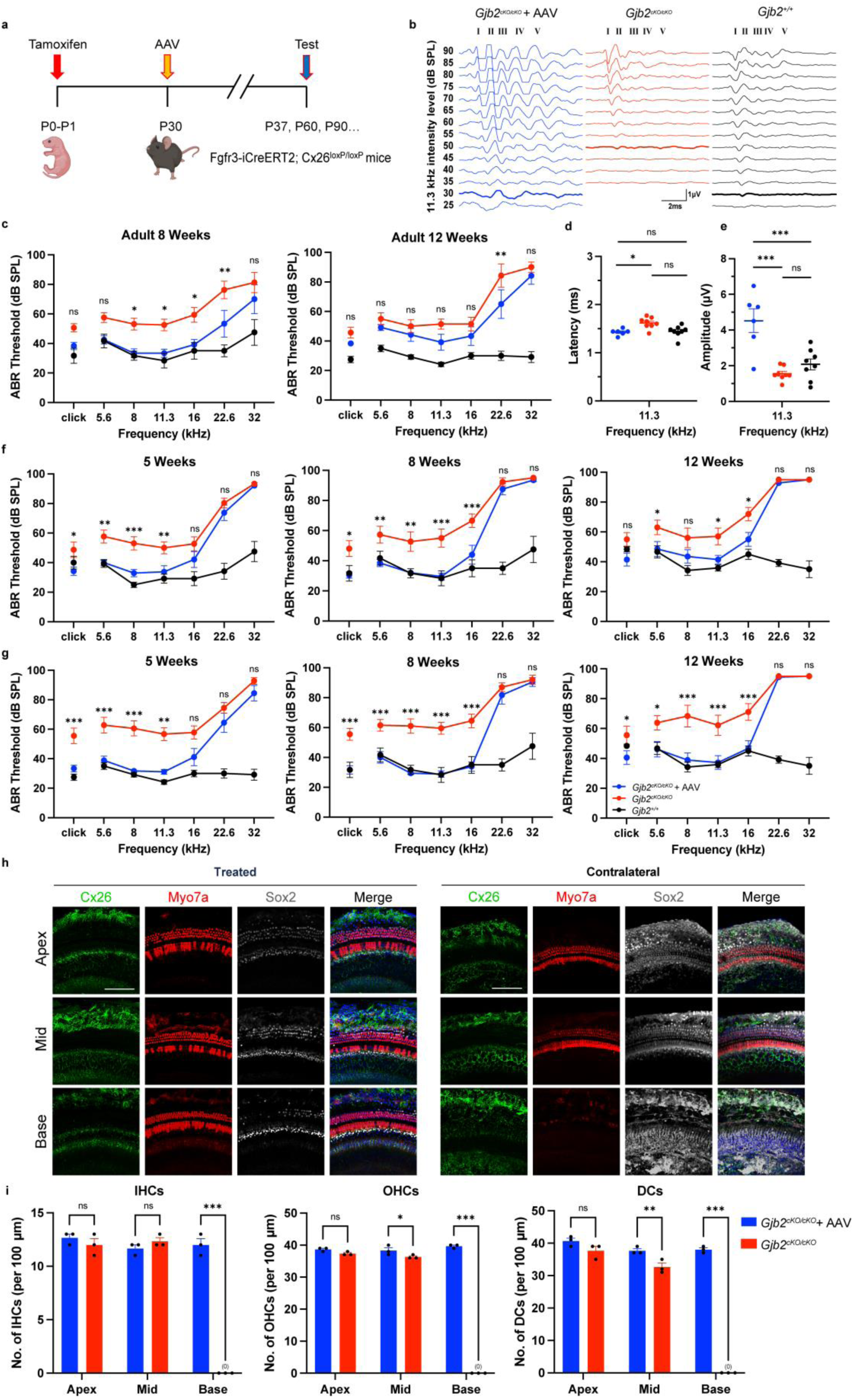
The auditory function of adult Fgfr3-iCreERT2; Cx26^loxP/loxP^ mice was rescued by administration of AAV-MAS012-WFS1-2274-mGJB2 or AAV-MAS012-WFS1-2274-hGJB2. **a,** Experimental design. **b,** Representative ABR traces in response to broadband click sound stimuli were recorded 1 week after AAV-MAS012-WFS1-2274-mGJB2 (titer: 2E13 VG/mL) administration at P30 Fgfr3-iCreERT2; Cx26^loxP/loxP^ mice. **c,** The ABR thresholds of Fgfr3-iCreERT2; Cx26^loxP/loxP^ mice for click sound stimuli and pure-tone stimuli were recorded 4 weeks (WT, n = 6; injected Fgfr3-iCreERT2; Cx26^loxP/loxP^ mice, n = 6; contralateral ear, n = 7) and 8 weeks (WT, n = 6; injected Fgfr3-iCreERT2; Cx26^loxP/loxP^ mice, n = 6; contralateral ear, n = 7) after AAV injection at P30. **d,** The ABR wave I latency in Fgfr3-iCreERT2; Cx26^loxP/loxP^ mice were recorded at 90 dB click sound intensities at 4 weeks after therapeutic injection at P30 (tone-burst stimuli at 90 dB and 11.3 kHz; WT, 1.44 ± 0.04 ms; injected ears, 1.42 ± 0.03 ms; *P* > 0.05) (WT, n = 6; injected Fgfr3-iCreERT2; Cx26^loxP/loxP^ mice, n = 6; contralateral ear, n = 7). **e,** The ABR wave I amplitude in Fgfr3-iCreERT2; Cx26^loxP/loxP^ mice were recorded at 90 dB click sound intensities at 4 weeks after therapeutic injection at P30 (sound intensity at 90 dB; WT, 2.07 ± 0.30 μV; injected ears, 4.51 ± 0.67 μV; *P* < 0.01) (WT, n = 6; injected Fgfr3-iCreERT2; Cx26^loxP/loxP^ mice, n = 6; contralateral ear, n = 7). **f,** The ABR thresholds of Fgfr3-iCreERT2; Cx26^loxP/loxP^ mice for click sound stimuli and pure-tone stimuli were recorded at 5 weeks, 8 weeks and 12 weeks of age after low dose (titer: 3×10^13^ VG/mL) AAV-MAS012-WFS1-2274-hGJB2 injection on P30 (WT, n = 6; injected Fgfr3-iCreERT2; Cx26^loxP/loxP^ mice, n = 12; contralateral ear, n=15). **g,** The ABR thresholds of Fgfr3-iCreERT2; Cx26^loxP/loxP^ mice for click sound stimuli and pure-tone stimuli were recorded at 5 weeks, 8 weeks and 12 weeks of age after high dose (titer: 5×10^13^ VG/mL) AAV-MAS012-WFS1-2274-hGJB2 injection on P30 (WT, n = 6; injected Fgfr3-iCreERT2; Cx26^loxP/loxP^ mice, n = 9; contralateral ear, n=9). **h,** AAV injection restored the expression of human Cx26 in PCs and DCs of the inner ear in Fgfr3-iCreERT2; Cx26^loxP/loxP^ mice. **i,** AAV injection inhibited the cell death of IHCs, OHCs and DCs in Fgfr3-iCreERT2; Cx26^loxP/loxP^ mice (treated Fgfr3-iCreERT2; Cx26^loxP/loxP^, *n* = 3; contralateral ear, n=3). Green: Cx26; Red: Myo7a; Gray: Sox2; Blue: DAPI. Scale bars: 100 μm.

Our ideal promoters should satisfy the following criteria: 1, The promoter sequences should enable specific and efficient transgene expression in inner ear cells that natively express Cx26. 2, The promoter sequences should avoid transgene expression in HCs. 3, The promoter sequences should also avoid transgene expression in other important organs, such as the central brain. Two weeks after inner ear injection of these AAVs through the round window membrane (RWM) into the right cochlea of neonatal (postnatal day 0-2, P0-P2) WT mice, efficiency and cell type-specific target range of each promoter were evaluated according to the expression pattern of fluorescent marker. A total of 5 out of the 21 tested synthetic promoters successfully drove the expression of the fluorescent marker in the inner ear (Extended Data Fig. 2 and Supplementary Table 2). Among these 5 promoters, the GJB2-1 promoter (AAV-ie-GJB2-1-EGFP) and WFS1-2274 promoter (AAV-ie-WFS1-2274-EGFP) fulfilled our screening criteria. In the AAV-ie-CMV-EGFP group, GFP was expressed in HCs, SCs, etc., (Fig. 1b). In contrast, the GJB2-1 promoter specifically restricted the expression of GFP to inner pillar cells (IPCs), outer pillar cells (OPCs), and DCs (Fig. 1b,c and Extended Data Fig. 3). Whole-mount immunofluorescence staining of the cochlea showed that the WFS1-2274 promoter specifically restricted the expression of GFP to Hensen’s cells (HeCs), DCs, OPCs, IPCs, and inner phalangeal cells (IPhCs) (Fig. 1b,c and Extended Data Fig. 3). Cryosectioning of the cochlea yielded results that were consistent with whole-mount immunofluorescence staining of the cochlea (Fig. 1d). Importantly, driven by either of these two promoters, GFP was not expressed in HCs (Fig. 1 b-d and Extended Data Fig. 3) or in the brains (Fig. 1e). Together these results indicate that our cell type-specific promoters exhibit strong cell type specificity.

### Cell type-specific promoter-mediated gene therapy provided long-term restoration of hearing in neonatal *Gjb2* cKO mice

During the period of pregnancy in mice, Cx26 is the major gap junction protein in gestational acinar cells, and these gap junctions are essential for normal development. Thus, mice with complete knockout of the *Gjb2* gene suffer from embryo death^28^. To develop a gene therapy system for *GJB2*-related genetic deafness, we constructed a *Gjb2* cKO mouse model by crossing Fgfr3-iCreERT2 mice with Cx26^loxP/loxP^ mice carrying the floxed *Gjb2* gene to obtain Fgfr3-iCreERT2; Cx26^loxP/loxP^ mice. Tamoxifen (TMX) injection into P0–P1 mice induced the Cre recombinase leading to deletion of Cx26 in DCs and PCs (hereafter the Fgfr3-iCreERT2; Cx26^loxP/loxP^ mice with TMX injection are referred to as *Gjb2* cKO mice)^30, 41^. At 4 weeks after injection of TMX, the *Gjb2* cKO mice exhibited significant hearing loss (Fig. 2), and at 8 weeks after injection, the *Gjb2* cKO mice exhibited significant knockout of Cx26 in DCs and PCs, whiles HCs and DCs were degenerated (Fig. 2). All of these results were consistent with previous research^30, 41^.

Several previous studies were able to partially restore the inner ear structural deficits caused by mutations of the *Gjb2* gene and to restore hearing function in newborn *Gjb2*-deficient mice^14, 28–30^. To evaluate the efficiency of our AAV-mediated *GJB2* gene therapy systems driven by the GJB2-1 and WFS1-2274 promoters on hearing recovery, we constructed plasmid vectors for gene replacement that expressed mouse *Gjb2* cDNA under the control of either the GJB2-1 or the WFS1-2274 promoter and packaged these two plasmid vectors into AAV-ie (AAV-ie-GJB2-1-mGJB2 and AAV-ie-WFS1-2274-mGJB2). ABR thresholds were measured to identify whether administration of AAV-ie-GJB2-1-mGJB2 or AAV-ie-WFS1-2274-mGJB2 could rescue the hearing of *Gjb2* cKO mice. Although injection of AAV-ie-WFS1-2274-mGJB2 significantly improved hearing in treated newborn *Gjb2* cKO mice (Extended Data Fig. 4), administration of AAV-ie-GJB2-1-mGJB2 more effectively restored auditory function in the treated newborn *Gjb2* cKO mice (Fig. 2a). Thus the efficiency of AAV-ie-GJB2-1-mGJB2 in restoring auditory function in newborn *Gjb2* cKO mice was further characterized. ABR thresholds were tested 4 weeks after injection of AAV-ie-GJB2-1-mGJB2 in *Gjb2* cKO mice. The click ABR thresholds of *Gjb2* cKO mice were 50.56 ± 4.3 dB, which were about 23.1 dB higher than the WT level (Fig. 2a), while the click ABR thresholds of AAV-ie-GJB2-1-mGJB2–treated mice were 34.33 ± 2.0 dB, which were about 16.2 dB lower than the *Gjb2* cKO mice (Fig. 2a). At 8 and 12 weeks post-injection, hearing was restored to WT levels for click, 5.6, 8, 11.3, and 16 kHz stimuli, whereas only partial hearing restoration was achieved at 22.6 and 32 kHz (Fig. 2a).

More importantly, we observed the long-term efficacy of AAV-ie-GJB2-1-mGJB2 gene therapy system, and at 16 and 26 weeks the hearing of treated *Gjb2* cKO mice was significantly improved to WT levels for click and 5.6, 8, 11.3, and 16 kHz stimuli (Fig. 2a). At 45 weeks after AAV injection, the hearing of the treated ear was restored to WT levels for click and 5.6, 8, and 11.3 kHz stimuli (Fig. 2a). These results indicate long-term effects of therapeutic efficacy and, to our knowledge, the longest-lasting hearing rescue reported for *GJB2*-associated hearing loss. Furthermore, we evaluated the safety of the AAV-ie-GJB2-1-mGJB2 therapeutic system. A total of 2 μL of therapeutic AAV was injected into the right ear of newborn WT mice, no significant hearing loss was observed at 4, 8, and 12 weeks post-injection (Extended Data Fig. 5), suggesting that the AAV-ie-GJB2-1-mGJB2 therapeutic system was well tolerated.

To test the efficiency of AAV-ie-GJB2-1-mGJB2 in re-expressing Cx26 protein and its efficiency in maintaining the integrity of the inner ear structure, we injected 2 μl of AAV into the right ears of newborn *Gjb2* cKO mice. Eight weeks after injection, Cx26 protein was successfully expressed in IPCs and OPCs of *Gjb2* cKO mice, while in the ears of *Gjb2* cKO mice without AAV injection, no Cx26 was found in PCs or DCs in the apical–middle–basal turns of the cochlea (Fig. 2b and Extended Data Fig. 6). More importantly, our results and previously published results demonstrated that *Gjb2* cKO mice exhibit HC and DC loss at P60 (Fig. 2b,c and Extended Data Fig. 6)^41^, HC counts showed the loss of almost all of the IHCs and OHCs in the basal turn, and DC counts showed the loss of almost all DCs in the basal turn (Fig. 2c and Extended Data Fig. 6). In AAV-injected *Gjb2* cKO mice, the HC and DC loss was alleviated (Fig. 2b,c and Extended Data Fig. 6). These results demonstrate that our gene therapy system could specifically restore the expression of Cx26 in IPCs and OPCs while protecting HCs and SCs from degeneration due to CX26 deficiency.

### Engineering of a novel AAV capsid to improve transduction efficiency in adult inner ear cells

Auditory function restoration in adult stage holds greater translational relevance for human clinical applications compared to interventions administered during the neonatal developmental stage. Our initial attempts to restore auditory function in adult *Gjb2* cKO mice using AAV-ie-GJB2-1-mGJB2 failed (Extended Data Fig. 7), and we attributed this phenomenon to the insufficient transduction efficiency of the AAV-ie in mature cochlear cells^32, 42^. To obtain capsids with higher gene delivery efficiency in mature cochlea, we initially investigated potential AAV receptors in the adult cochlea cells and discovered that integrins are broadly expressed within the cochlea. Specifically, we observed that SCs, including PCs, DCs, inner border phalangeal cells, HeCs, as well as stria vascularis (SV) cells, such as those in intermediate stria and marginal stria, expressed high levels of genes encoding RGD (Arg-Gly-Asp) integrins subunits, including ITGAV, ITGA5, ITGB1, ITGB3 and ITGB5 (Extended Data Fig. 8)^43^. Peptides containing the RGD motif designed to target integrins can significantly enhance the transduction of AAV9 capsids in muscle tissue^44^. Therefore, we selected one of these peptides, myo2A (GPGRGDQTTL), and inserted it into the AAV-DJ capsid between N589 and R590, resulting in the creation of a new capsid variant, MAS012 (Fig. 3a). To test the transduction efficiency of AAV-MAS012 in adult inner ear cells, we packaged AAV-MAS012-CAG-nls-tdTomato and AAV-ie-CAG-nls-tdTomato, and at two weeks after injection of AAVs into the inner ear of P30 WT mice, we measured the expression range of tdTomato. Compared with the AAV-ie group, the AAV-MAS012 group showed increased transduction efficiency in mature SCs, while the transduction efficiency in HCs was limited (Fig. 3b,c).

To further validate the transduction efficiency of AAV-MAS012 in mature cochlear cells, we packaged AAV-MAS012-WFS1-2274-EGFP and AAV-ie-WFS1-2274-EGFP, because the WFS1-2274 promoter worked in a larger target range than the GJB2-1 promoter. Two weeks after injection, the whole-mount immunofluorescence staining results showed that GFP was mainly expressed in HeCs, DCs, OPCs, IPCs, and IPhCs in the AAV-MAS012 group, while the transduction efficiency of AAV-ie was lower, and GFP was not expressed in HCs in either groups (Fig. 3d,e). Cryosection analysis of cochleae treated with AAV-MAS012-WFS1-2274-EGFP yielded results that were consistent with the whole-mount immunofluorescence staining that GFP was expressed in SCs, besides, GFP was also expressed in the SV, the SPL and the SL (Fig. 3f). These findings indicate a greater transduction efficiency for AAV-MAS012 compared to AAV-ie and show that GFP expression driven by the WFS1-2274 promoter closely matches the expression pattern of Cx26 protein in the inner ear.

### The *GJB2* gene therapy system using AAV-MAS012 and cell type-specific promoters restored hearing in adult *Gjb2* cKO mice

Before we tested whether AAV-MAS012 could be used to treat the hearing loss in adult *Gjb2* cKO mice, both AAV-MAS012-GJB2-1-mGJB2 and AAV-MAS012-WFS1-2274-mGJB were used to rescue the auditory function of *Gjb2* cKO mice after the onset of hearing (P11–P14)^2, 45^. Unilaterally injection 2 µL of GJB2-1-mGJB2 or WFS1-2274-mGJB2 at P14 could restored the hearing function of the *Gjb2* cKO mice (Extended Data Fig. 9), with administration of AAV-MAS012-WFS1-2274-mGJB2 restoring hearing to WT levels for clicks and for 5.6, 8, 11.3, and 16 kHz stimuli (Extended Data Fig. 9). So, we chose the AAV-MAS012-WFS1-2274-mGJB2 system for the treatment of adult *Gjb2* cKO mice.

We unilaterally injected AAV-MAS012-WFS1-2274-mGJB2 into the inner ear of *Gjb2* cKO mice at P30 (Fig. 4a). Compared with untreated mice, at four weeks after AAV administration we observed significantly decreased ABR thresholds almost to WT levels for click and 5.6, 8, 11.3, and 16 kHz stimuli, while the hearing restoration at 22.6 and 32 kHz was inferior to WT levels (Fig. 4b,c). The ABR wave I latency was similar to WT levels (Fig. 4d), while the wave I amplitudes in the ABR waveforms at 11.3 kHz were higher than WT (Fig. 4e). More importantly, hearing was restored in adult *Gjb2* cKO mice at 8 weeks after AAV administration (Fig. 4c). Together these results demonstrated that cell type-specific promoters delivered by the high transduction efficiency AAV-MAS012 could restore hearing function in adult *Gjb2* cKO mice.

### Improvement of hearing function using human *GJB2* cDNA

To further facilitate the clinical translation of gene therapy for *GJB2*-related hereditary deafness, we replaced the mouse *Gjb2* cDNA with human *GJB2* cDNA to create AAV-MAS012-WFS1-2274-hGJB2. At 1 and 4 weeks after injection of a low-dose of AAV into P30 *Gjb2* cKO mice, hearing function was restored almost to WT levels across frequencies from clicks to 16 kHz. This restoration of hearing function was sustained to at least 12 weeks of age (Fig. 4f). In the high-dose AAV injection group, hearing function was restored to WT levels across frequencies from clicks to 16 kHz at 5 weeks, 8 weeks, and 12 weeks of age (Fig. 5g). Consistent with hearing restoration, AAV injection restored Cx26 expression in the SCs and prevented the loss of IHCs, OHCs, and DCs (Fig. 4h,i). Taken together, these results suggest that the *GJB2* gene therapy system we constructed, which consists of the newly constructed AAV-MAS012 capsid and the cell type-specific WFS1-2274 promoter, has great potential for clinical application.

**Fig. 5.**
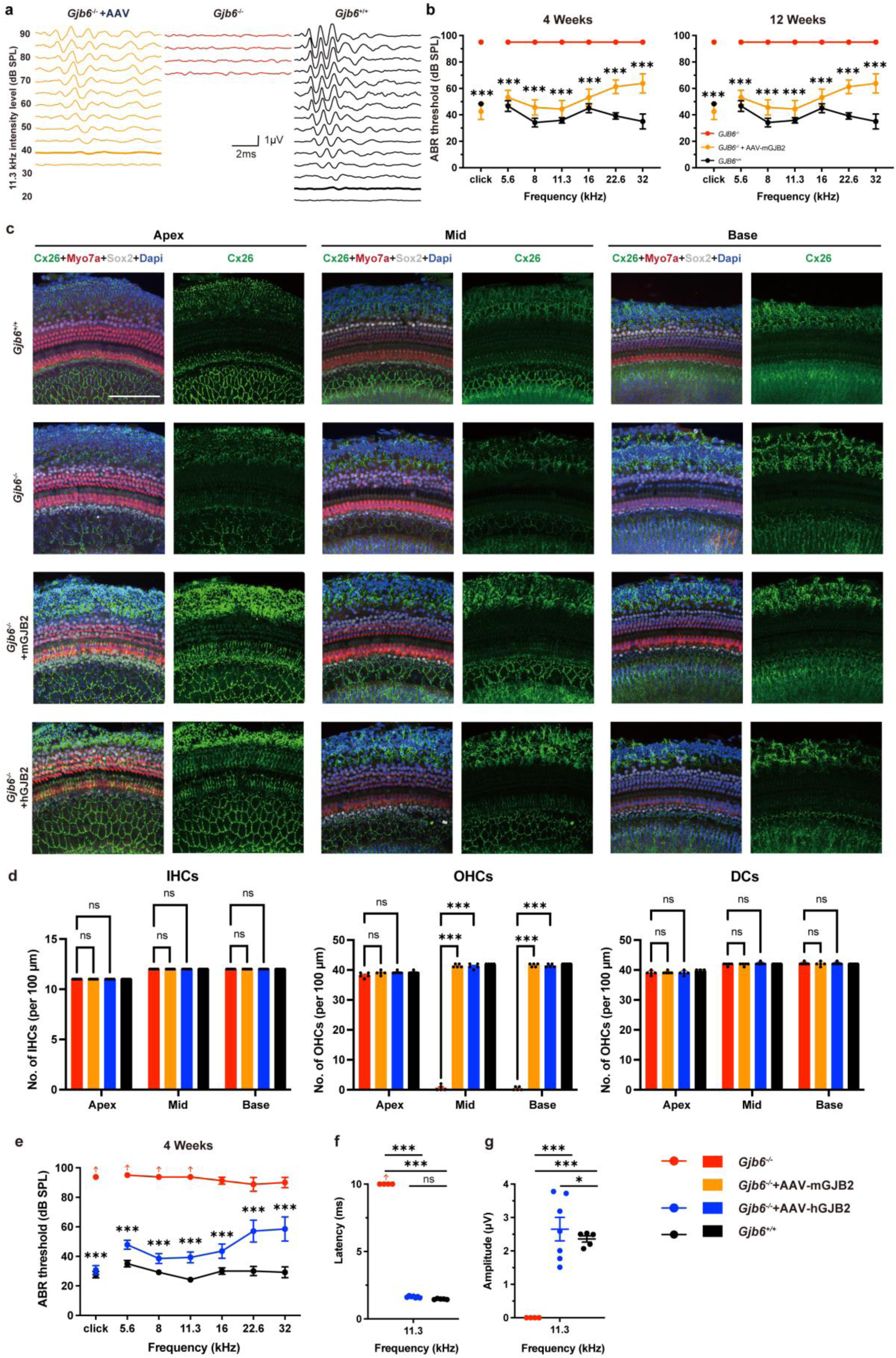
AAV-mediated gene therapy restored auditory function in different *Gjb2*-deficient mouse models. **a,** Representative ABR traces in response to broadband click sound stimuli after AAV-MAS012-WFS1-2274-mGJB2 (titer: 3E13 VG/mL) administration in newborn *Gjb6^−/−^*mice. **b,** The ABR thresholds of *Gjb6^−/−^*mice for click sound stimuli and pure-tone stimuli were recorded at 4 weeks and 8 weeks after AAV injection at P0–P2 (WT, n = 6; *Gjb6^−/−^* mice, n = 4; injected *Gjb6^−/−^* mice, n = 8). **c,** AAV injection restored the expression of mouse Cx26 in PCs and DCs of the inner ear in *Gjb6^−/−^* mice and inhibited cell death in the inner ear. **d,** Quantification of the inhibition of cell death of IHCs, OHCs, and DCs after AAV injection in *Gjb6^−/−^* mice (*Gjb6^−/−^* mice, n = 5; AAV-mGJB2 treated *Gjb6^−/−^* mice, n = 5; AAV-hGJB2 treated *Gjb6^−/−^* mice, n = 5; WT, n = 5). **e,** The ABR thresholds of *Gjb6^−/−^* mice for click sound stimuli and pure-tone stimuli were recorded at 4 weeks after AAV-MAS012-WFS1-2274-hGJB2 (titer: 7E13 VG/mL) injection (*Gjb6^−/−^*mice, n = 4; injected *Gjb6^−/−^*, n = 7; WT, n = 6). **f,** The ABR wave I latencies in *Gjb6^−/−^* mice were recorded at 90 dB click sound intensities 4 weeks after AAV-MAS012-WFS1-2274-hGJB2 injection (tone-burst stimuli at 90 dB at 11.3 kHz; WT, 1.47 ± 0.02 ms; injected ears, 1.65 ± 0.03 ms; *P* > 0.05) (*Gjb6^−/−^* mice, n = 4; injected *Gjb6^−/−^*, n = 7; WT, n = 6). **g,** The ABR wave I amplitudes in *Gjb6^−/−^*mice were recorded at 90 dB click sound intensities 4 weeks after AAV-MAS012-WFS1-2274-hGJB2 injection (sound intensity at 90 dB; WT, 2.36 ± 0.09 μV; injected ears, 2.32 ± 0.39 μV; *P* < 0.05) (*Gjb6^−/−^*mice, n = 4; injected *Gjb6^− /−^*, n = 7; WT, n = 6). Green: Cx26; Red: Myo7a; Gray: Sox2; Blue: DAPI. Scale bars: 100 μm.

To visualize exogenous Cx26 expression, 3**×**hemagglutinin (HA) tag was added to the C-terminal of hGJB2 in the therapeutic system. One week after AAV injection at P30, the hearing of treated *Gjb2* cKO mice was restored (Extended Data Fig. 10a). At 8 weeks after AAV administration, immunofluorescence results showed that exogenous Cx26 was expressed in the HeCs, DCs, OPCs, IPCs, IPhCs, and HCs of the injected ear, but not in the contralateral ear (Extended Data Fig. 10b). These results are largely consistent with those obtained using AAV-MAS012-WFS1-2274-EGFP (Fig. 3d-f), and all of these results suggest that the expression of the transgenes delivered by our gene therapy system effectively cover the expression range of Cx26 itself within the inner ear.

### Confirming the universality of cell type-specific promoters in the *GJB2* gene therapy system

To determine whether AAV-MAS012 delivery of cell type-specific promoters in the *GJB2* gene therapy system could also restore the hearing function of other *Gjb2* mutant mouse models, we injected AAV-MAS012-WFS1-2274-mGJB2 into the inner ear of newborn *Gjb6* null mutant mice (*Gjb6*^LacZ/LacZ^, hereafter referred to as *Gjb6*^−/−^)^29, 46–48^. *Gjb6^−/−^* mice are another animal model of DFNB1 and are characterized by severe congenital nonsyndromic deafness^46, 49^. Four weeks after injection of AAV-MAS012-WFS1-2274-mGJB2 at P0-P2, the hearing function of the injected ears was significantly improved (Fig. 5a,b), and the restoration of hearing could be maintained to at least 12 weeks (Fig. 5b). The expression of Cx26 was apparently restored in SCs (Fig. 5c), while the OHC loss was significantly inhibited after AAV administration (Fig. 5c,d).

To further validate the efficacy of our humanized *GJB2* gene therapy system, 2 µL of AAV-MAS012-WFS1-2274-hGJB2 was injected into the right ear of newbore *Gjb6^−/−^* mice. Four weeks post-injection, hearing thresholds in the injected ears showed marked restoration across all frequencies, and click ABR thresholds were restored to approximately 35 dB—around 58.75 dB lower than the thresholds in the untreated *Gjb6^−/−^* mice. Under tone-burst stimuli, the average hearing thresholds in the injected ears were significantly improved at click and 5.6, 8, 11.3, and 16 kHz compared with untreated controls (Fig. 5e). The ABR wave I latency was similar to WT levels (Fig. 5f), while the wave I amplitudes in the ABR waveforms at 11.3 kHz were higher than WT (Fig. 5g). The expression of Cx26 protein was also restored in SCs (Fig. 5c), and the HCs loss was inhibited after AAV injection (Fig. 5d). Together these results indicate that our humanized *GJB2* gene therapy system can ameliorate hearing loss across different types of *Gjb2*-deficient mouse models.

### Safety and transduction assessment of humanized *GJB2* gene therapy system in non-human primate

Non-human primates (NHPs) represent a more suitable animal model for evaluating transgenes delivery to the human inner ear^28, 50^. To advance our *GJB2* gene therapy system toward clinical application, we evaluated auditory toxicity and systemic toxicity after local administration of AAV-MAS012-WFS1-2274-hGJB2 in cynomolgus monkeys. Two monkeys received bilateral low-dose injections of AAV-MAS012-WFS1-2274-hGJB2 and were monitored for auditory toxicity until 4 weeks post-injection, after which they were euthanized for systemic toxicity assessment. One monkey received bilateral high-dose AAV injection and was monitored for auditory safety up to 10 weeks post-injection. All animals exhibited normal auditory function prior to AAV administration (Fig. 6a,b). The AAV was administered into the cochleae via the RWM. Compared to preoperative baseline levels, some injected ears exhibited elevated ABR thresholds one-week post-injection. However, by four weeks in both low- and high-dose groups, and by ten weeks in the high-dose group, ABR thresholds recovered to near-normal levels. (Fig. 6a,b). These findings indicated that AAV injection does not impair the normal auditory function in cynomolgus monkeys and induced no significant changes in haematological, coagulation or serum biochemistry parameters relative to pre-injection baselines (Supplementary Table 3).

**Fig. 6.**
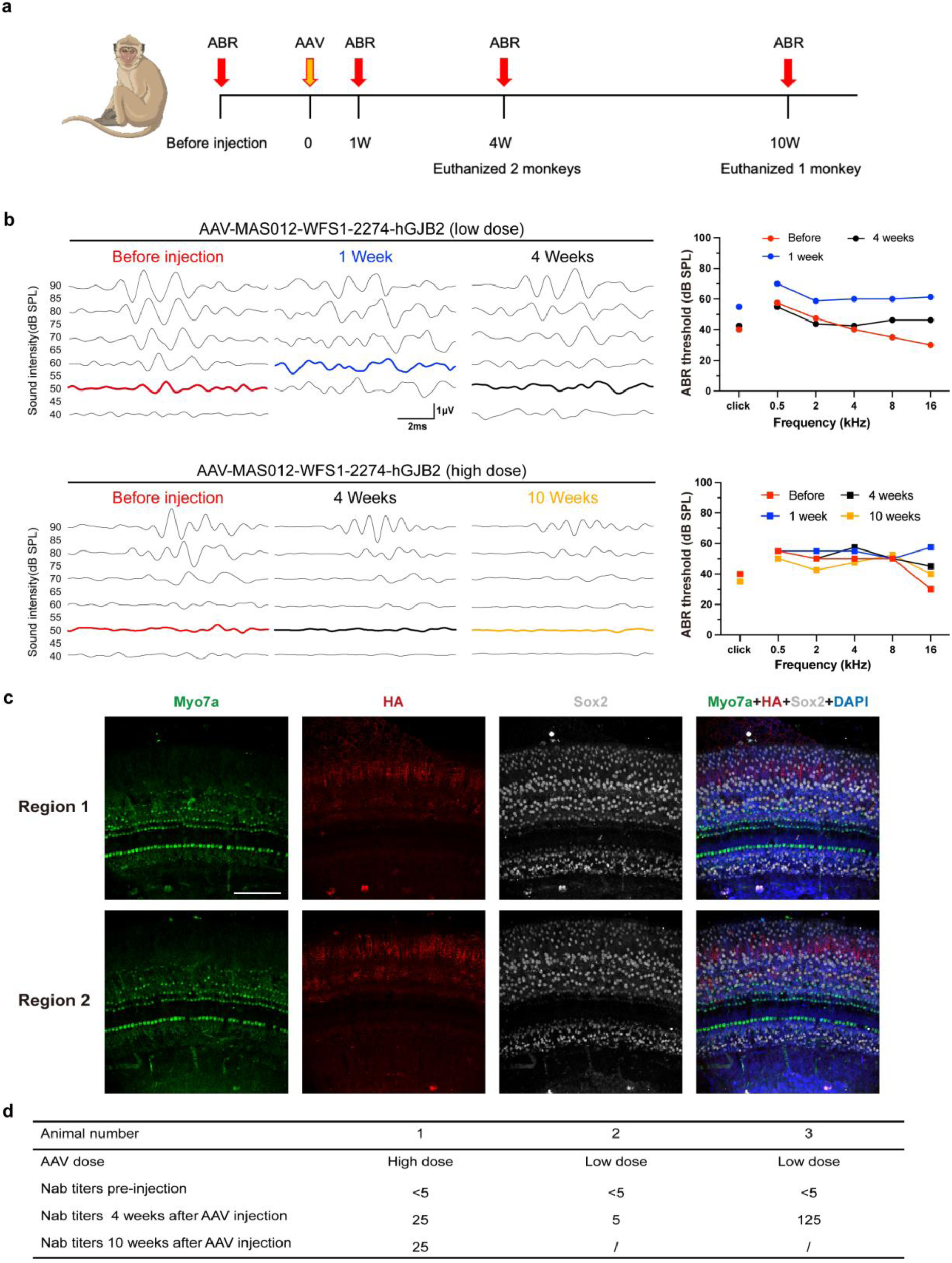
Evaluation of the safety and transduction efficacy of *GJB2* gene therapy system in nonhuman primates. **a,** Schema of the experimental procedure. **b,** Representative ABR traces in response to broadband click sound stimuli before and after AAV-MAS012-WFS1-2274-hGJB2 administration in cynomolgus monkeys. **c,** The ABR thresholds of cynomolgus monkeys for click sound stimuli and pure-tone stimuli were recorded before AAV injection and at 1 week, 4 weeks, and 10 weeks after AAV injection (low dose titer: 2.7E13 VG/mL, 2 monkeys, 4 ears; high dose titer: 7E13 VG/mL, 1 monkey, 2 ears). **d,** The immunofluorescence results of the HA tag showed the expression pattern of exogenous Cx26 in the AAV-MAS012-WFS1-2274-hGJB2-3×HA (titer: 3.9E13 VG/mL) injected ear of cynomolgus monkey. **e,** NAb titers in serum before and after AAV-MAS012-WFS1-2274-hGJB2 administration. Green: Cx26; Red: HA-tag; Gray: Sox2; Blue: DAPI. Scale bars: 100 μm.

We also evaluated the transduction efficiency of the *GJB2* gene therapy system in cynomolgus monkey. To identify the exogenous *GJB2* expression in the inner ear, we injected AAV-MAS012-WFS1-2274-hGJB2-3×HA carrying a C-terminal HA tag into the inner ear of a cynomolgus monkey bilaterally. Immunofluorescence analysis confirmed efficient expression of exogenous Cx26 in the injected ear (Fig. 6c), indicating that our therapeutic system performs sufficient transduction efficiency in mature cynomolgus monkey inner ear SCs. We also identify the serum neutralizing antibodies (NAbs) before and after AAV injection. NAb titers at baseline in all cynomolgus monkeys were <1:5, four weeks and 10 weeks after AAV injection, the serum NAb titer increased to 1:25 in high dose AAV-MAS012-WFS1-2274-hGJB2 group, while the serum NAb titer were 1:5 and 1:125 in low dose group four weeks after AAV administration (Fig. 6d). The administration of AAV-MAS012-WFS1-2274-hGJB2 appears to result in a low to moderate immunogenicity response in cynomolgus monkeys, further confirming the safety profile of our therapeutic system. Taken together, these results indicated that our humanized *GJB2* gene therapy system exhibits favorable safety and transduction efficiency, as well as acceptable immunogenicity in non-human primates.

## Discussion

One of the key obstacles limiting the development of AAV-mediated *GJB2* gene therapy drugs is that ectopic expression of Cx26 in the inner ear induces HC apoptosis and leads to hearing dysfunction^12, 28^. In this study, in order to specifically deliver *GJB2* cDNA into cells of the inner ear that natively express Cx26, we designed a library of 21 different synthetic promoter sequences that were constructed by four different strategies. Among these 21 different promoters, the GJB2-1 promoter and WFS1-2274 promoter met our requirements. The GJB2-1 promoter restricted the expression of GFP to IPCs, OPCs, and DCs, and the WFS1-2274 promoter restricted the expression of GFP to HeCs, DCs, IPCs, OPCs, and IPhCs, as well as to the SV, the SPL and the SL. We validated the efficiency of these two promoters by driving the expression of Cx26 protein. Controlled by the GJB2-1 promoter, the expression of Cx26 protein in the DCs, IPCs, and OPCs of *Gjb2* cKO mice was restored, the loss of OHCs and DCs was inhibited, and the hearing function was rescued. In addition, at 45 weeks after AAV injection the hearing function of treated *Gjb2* cKO mice was markedly improved. To our knowledge, this is the longest-lasting treatment system among all reported *GJB2*-related genetic deafness gene therapy drugs. The promoter is an important component of an expression vector because it controls when and where a transgene is expressed in an organism. Many recently published gene therapy drugs for hereditary deafness have made use of cell type-specific promoters^4, 28–30^. Given that efficient targeted delivery of transgenes into GJB2-expressing SCs using AAVs remains a major obstacle, it is particularly important to generate cell type-specific promoters to restrict the expression range of Cx26 to specific cell populations^51, 52^.

Another key challenge hindering the development of AAV-mediated *GJB2* gene therapy drugs is the relatively low transduction efficiency of AAV in mature inner ear cells. In this study, AAV-ie–mediated gene therapy was unable to restore hearing function in adult *Gjb2* cKO mice (Extended Data Fig. **7**). By inserting an RGD peptide-containing fragment into the AAV-DJ capsid, we successfully engineered a novel AAV-MAS012 capsid that exhibits high transduction efficiency in mature inner ear SCs. In fact, beyond transducing SCs and the SV, AAV-MAS012 exhibits a broader tropism within the inner ear. Highly overlapping with endogenous Cx26 expression, AAV-MAS012 also effectively transduces the SPL and the SL. This suggests that integrins may also be expressed in these regions, thereby facilitating AAV transduction, which is worth further exploring. This discovery will provide new ideas for the targeted modification of AAV capsids in the inner ear of adult mice. Delivered by AAV-MAS012, our cell type-specific WFS1-2274 promoter driving the *GJB2* gene therapy system successfully rescued the hearing of adult *Gjb2* cKO mice. This is the first time that *GJB2* gene therapy system has been successfully applied to restore the hearing function of adult *Gjb2*-deficient mice.

To develop a potential therapeutic system for treating human hearing loss caused by mutations in the *GJB2* gene, we constructed an AAV-based therapeutic system using the coding region of the human *GJB2* gene. The expression of human Cx26 in the inner ear restored hearing in adult Fgfr3-iCreERT2; Cx26^loxP/loxP^ mice to near WT levels for click stimuli and for tone-burst stimuli up to 16 kHz for at least 2 months. In addition to *Gjb2* cKO mice, our *GJB2* gene therapy systems also restored hearing function in *Gjb6^−/−^* mice. More importantly, the humanized *GJB2* gene therapy system demonstrates sufficient safety and transduction efficiency in non-human primates. All these results indicate that our GJB2 therapeutic system has broad therapeutic efficacy across various Cx26-deficient animal models, along with a favorable safety profile, thereby enhances its potential for clinical translation.

In summary, by developing cell type-specific promoters and engineering a novel high-efficiency AAV capsid that targets mature cells in the inner ear we successfully established a gene therapy platform for *GJB2*-related hereditary deafness. The safety and efficacy of our humanized *GJB2* gene therapy system was validated in different *Gjb2*-deficient mouse models and non-human primates, positioning it as a promising candidate for future clinical translation.

## Methods

### Animals

All animal care and all animal experiments were performed in accordance with the Ethics Committee of Fudan University, and the Shanghai Medical Experimental Animal Administrative Committee, China. Mice were housed in groups of six in ventilated, pathogen-free cages with free access to water and food. The temperature was about 25℃ and the dark/light cycle was 12 hours. Wild type (WT) C57BL/6J mice were injected with AAVs to evaluate the efficiency and cell-type specificity of the different promoters and the transduction efficiency of new AAV serotypes. The Cx26^loxP/loxP^ mouse line and the Fgfr3-iCreERT2 mouse line were provided by Prof. Xi Lin from Emory University, Atlanta, USA, and Prof. Zhiyong Liu of the Chinese Academy of Sciences, Shanghai, China, respectively. The two sexually mature mouse lines were crossbred to obtain Fgfr3-iCreERT2; Cx26^loxP/WT^ mice, and their offspring were intercrossed to generate Fgfr3-iCreERT2; Cx26^loxP/loxP^ double transgenic mice. Tamoxifen (TMX) (1.5 mg/10 g body weight) was injected subcutaneously into both male and female mice on the day of birth (P0) and the first day after birth (P1) to activate Cre recombinase and achieve targeted knockout of Cx26 in DCs and PCs in the inner ear of the Fgfr3-iCreERT2; Cx26^loxP/loxP^ mice^41^. PCR amplification of genomic DNA from tail tissues was used to determine the genotypes of the Fgfr3-iCreERT2; Cx26^loxP/loxP^ mice. The constitutive knockout *Gjb6^−/−^* mice were purchased from the European Mouse Mutant Archive (EMMA, #EM:00323, www.infrafrontier.eu) and have been used in several previously published studies^29, 41, 46^. Male and female mice were randomly selected for all experiments. The cynomolgus monkeys (two males and two females) were provided by TriApex Laboratories Co., Ltd (Nanjing, China). Three to six years old cynomolgus monkeys were used in this research, and all animals were housed in well-ventilated rearing cages, separately. The rearing temperature was about 19℃–26℃, humidity was about of 40%–70%, and the dark/light cycle was 12 hours.

### Design of the cell type-specific promoter library

Based on the RNA sequencing data and the possible positions of the promoter sequences in the gene, we developed the cell type-specific promoter library based on a rational design strategy and a large-scale screening strategy. First, we predicted two GJB2 promoter sequence locations in the *Gjb2* gene: a 3628 bp (–3628 bp to –1 bp) GJB2-1 promoter sequence and a 3600 kb (–7200 bp to –3601 bp) GJB2-2 promoter sequence upstream of the start codon of the mouse *Gjb2* gene. Second, based on single-cell RNA-seq data, three different DNA sequences in the *Gjb2* gene were predicted as candidate GJB2 promoter sequences (2001 bp, 900 bp, and 411 bp, respectively). Based on the sequence information, we constructed a 2001 bp Y1 promoter, a 2901 bp (2001 bp + 900 bp) Y2 promoter, and a 3312 bp (2001 bp +9 00 bp + 411 bp) Y3 promoter. To further increase the efficiency of these promoters, we added a 380 bp CMV enhancer to Y1 and Y3, generating a 2381 bp Y1E promoter and a 3592 bp (2001 bp + 900 bp + 411 bp + 380 bp) Y3E promoter. Third, in previous studies^35, 36^, we identified several GJB2 promoter candidate sequences of human origin along with a 1043 bp C3 enhancer that could increase the efficiency of the human-origin GJB2 promoters. We compared the sequences of these promoters, selected two promoter sequences and modified them by adding the C3 enhancer, creating a 2581 bp (1541 bp +1043 bp) hGJB2-2 promoter and a 3768 bp (2725 bp +1043 bp) hGJB2-10 promoter. Fourth, six genes (*Gjb6*, *Myh14*, *Wfs1*, *Eya1*, *Sox10*, and *Myh9*) were found to have similar expression patterns in the inner ear as *Gjb2* (Supplementary Table 1). Six promoters with a length of about 3450 bp (≈–3450 bp to –1 bp) upstream of the start codon of these genes (GJB6-3450 promoter, MYH14-3450 promoter, WFS1-3450 promoter, EYA1-3450 promoter, SOX10-3450 promoter, and MYH9-3450 promoter) and six promoters with a length of about 2000 bp (≈–2000 bp to –1 bp) upstream of the TSS of these genes (GJB6-2000 promoter, MYH14-2000 promoter, WFS1-2274 promoter, EYA1-2000 promoter, SOX10-2000 promoter, and MYH9-2000 promoter) were predicted^37, 38^. Using the above four methods, we obtained 21 cell type-specific promoter sequences. We constructed 21 AAV plasmids containing these promoters and packaged the recombinant plasmids into AAV-ie to drive the expression of a fluorescent marker. These AAVs were then injected into the inner ears of newborn WT mice. The relevant tissues (cochlear and central brain) were harvested for immunohistochemistry to identify the cell type specificity of these promoters.

### AAV plasmid construction

All cell type-specific promoter sequences were subcloned into pAAV-CMV-EGFP or pAAV-CMV-mNeonGreen-Bpnls to replace the cytomegalovirus (CMV) promoters, while CMV immediate-early enhancer/chicken β-actin (CAG) promoter and CMV promoter were used as ubiquitous promoter controls. For gene therapy in *Gjb2* mutant mice, the mouse *Gjb2* coding sequence (CDS) (NM_008125.3) or human *GJB2* CDS (NM_004004.6) was packaged into AAV-ie viruses or AAV-MAS012 (PackGene Biotech). The AAV vector plasmids contained a promoter (GJB2-1 or WFS1-2274), a poly-adenylation sequence, and a Kozak sequence.

### AAV production and purification

All AAVs were produced by PackGene Biotech or in our lab utilizing HEK 293T cells through a triple-transfection method as previously described^53^. The transfection reaction included pAAV plasmids carrying the gene of interest, pHelper, and the appropriate capsid plasmids, all in a molar ratio of 1:1:1. Cell pellets and the media were collected 5 days post-transfection. The viral particles from the cell pellets were released using a triple freeze/thaw cycle. The viral particles present in the media were isolated and concentrated through a precipitation process that involved the use of polyethylene glycol (PEG) in combination with sodium chloride (NaCl). rAAVs were then purified by iodixanol gradient centrifugation. The titer of the final AAV stock was determined by performing a quantitative polymerase chain reaction (qPCR) assay. The vector genome (VG) titers of AAV-ie-EGFP and AAV-ie-mNeonGreen-Bpnls were 1 × 10^13^ VG/mL (PackGene Biotech), the VG titers of AAV-ie-CAG-mGJB2, AAV-ie-GJB2-1-mGJB2, and AAV-ie-WFS1-2274-mGJB2 were 2 × 10^13^ VG/mL (PackGene Biotech), the VG titers of AAV-MAS012-GJB2-1-mGJB2 was 1 × 10^13^ VG/mL (packed by our lab), the VG titers of AAV-MAS012-WFS1-2274 -mGJB2 was 1 × 10^13^ VG/mL (packed by our lab) or 3 × 10^13^ VG/mL (PackGene Biotech), the VG titers of AAV-MAS012-WFS1-2274-hGJB2 was 2.7 × 10^13^ VG/mL, 3 × 10^13^ VG/mL, 5 × 10^13^ VG/mL or 7 × 10^13^ VG/ml (PackGene Biotech and our lab), the VG titers of AAV-MAS012-CAG-nls-tdTomato and AAV-ie-CAG-nls-tdTomato were 1 × 10^13^ VG/mL. To visualize exogenous Cx26 expression, the hemagglutinin (HA) tag was added to the C-terminus of the human GJB2 CDS, and the VG titers of AAV-MAS012-WFS1-2274 pro-hGJB2-3xHA was 3 × 10^13^ VG/mL or 3.9 × 10^13^ VG/mL (PackGene Biotech).

### AAV administration in mice

AAV was microinjected into the right ears of the newborn mice via the round window membrane (RWM) as we have previously described^9, 34^. The newborn P2-P3 mice were anesthetized by placing them on ice during the operation. The skin behind the right ear was incised to expose the RWM covered by the auditory bulla. A total of 2 µL of AAV-ie-EGFP, 2 µL of AAV-ie-mNeonGreen-Bpnls or 2 µL of AAV-ie-GJB2 therapeutic agents were injected into the inner ears with a glass micropipette (504949, World Precision Instruments) connected to a Nanoliter Microinjection System (NANOLITER2020, World Precision Instruments). The injection rate was set to 6 nL/s. The incision was closed after injection, and the pups were placed on a 37℃ heating pad for 10 minutes to fully recover. Adult mice were anesthetized with ketamine (60 mg/kg) and xylazine (60 mg/kg) through intraperitoneal injection. During the surgical procedure, the mice were placed on a 37℃ heating pad to maintain their body temperature. After the fur of the post-auricular region of the right ear was shaved and cleaned, an incision was made and the sternocleidomastoid muscle was transected to expose the posterior semicircular canal (PSCC), and a PSCC fenestration was performed with a 26-G needle. A glass micropipette was then connected to the above-mentioned Nanoliter Microinjection System and inserted into the PSCC. A total of 2 µL of AAV-GJB2 therapeutic agent was injected into the inner ear at 6 nL/s. After injection, the hole was sealed with tissue adhesive (1469SB, 3M, America), and the incision was closed after injection. The mice were placed on a 37℃ heating pad for full recovery.

### AAV administration in cynomolgus monkeys

AAV was microinjected into the both ears of cynomolgus monkeys via the RWM route as previously described^28, 50^. The cynomolgus monkeys were general anesthetized, with heart rate, respiratory rate, and oxygen saturation monitored throughout the surgical procedure. A 25-millimeter semilunar incision was made to expose the mastoid cortex. Fine hemostasis was achieved intraoperatively using a bipolar electrocautery device. The external auditory canal wall, tympanic membrane, facial nerve, and chorda tympani nerve were carefully protected. After exposing the round window niche and the stapes footplate, a fenestration was created at the oval window using a needle. The site was observed for several minutes until no significant perilymph leakage was noted. The bony overhang of the round window niche was gently removed to fully expose the RWM. A nano-liter scale microinjection system connected to an injection needle was used to continuously infuse approximately 50 µL of AAV into each ear at a rate of 2-3 µL/min via the round window membrane. After the injection, muscle tissue was used to seal both the RWM and the fenestration on the stapes footplate. The mastoid cavity was filled and closed with bone powder and adjacent autologous tissue, and the skin was sutured closed.

### Venous blood and cerebrospinal fluid sampling in cynomolgus monkeys

The protocols were performed according to previous study^50^. On postoperative days 7、28 and 56, blood was drawn intravenously from the femoral or saphenous veins in anesthetized animals after disinfecting the epidermis with 75% alcohol. A standard vacuum blood collection needle was inserted into the vein, and after withdrawal, a cotton swab was applied to compress the vessel for hemostasis. Serum biochemical and routine blood tests for non-human primates were performed by TriApex Laboratories Co., Ltd. (Nanjing, China). Prior to euthanasia, cerebrospinal fluid was collected from the cisterna magna after preoperative administration of atropine. Following the induction of general anesthesia, the hair over the cisterna magna was removed and the skin was disinfected with 75% alcohol. The animal’s neck was then flexed to adequately expose the area, and a 23G needle was used to collect the cerebrospinal fluid sample.

### Immunohistochemistry and confocal microscopy

To identify the strength and cell type-specificity of the different promoters, cochlear whole mounts of injected ears (P30) and un-injected ears were immunostained as previously described in mice^9, 33^. The temporal bone of the dissected cochleae was perforated, and the cochlea was perfused and fixed with 4% fresh paraformaldehyde (PFA) solution at room temperature for 1–2 h and decalcified with 10% ethylene diamine tetraacetic acid (EDTA) for 4–6 h at room temperature. Mouse brains were fixed in 4% fresh PFA solution at 4°C for 24–48 h and sectioned with a vibrating blade microtome (Leica) at a thickness of 50 μm. For immunohistochemical experiments in non-human primates, all animals were euthanized under deep anesthesia without recovery as previous described^50^. Specifically, after exposing the carotid artery, thorough perfusion was first performed with saline, followed by injection of formalin solution to achieve complete flushing and preliminary fixation of the inner ear samples. Subsequently, cochlear tissue samples were collected for histological studies. The cochleae were fixed with 4% PFA solution and decalcified with 10% EDTA for about 1 month at room temperature. Then the basilar membrane was collected from the cochlea. The tissues were washed three times with PBS, permeabilized in 0.02% Triton X-100 for 30 min at room temperature, and blocked with PBS containing 1% Triton X-100 (1% PBST) and 10% donkey serum for approximately 14h at 4°C. The tissues were then incubated with the primary antibody overnight at 4°C. The following primary antibodies were used in this study: mouse anti-GJB2 antibody (1:200 dilution, Invitrogen, 13-8100) was used as the indicator of GJB2 expression; rabbit anti-Myosin-VIIa (1:400 dilution, Proteus BioSciences, 25-6790) was used as the indicator of HCs; goat anti-Sox10 (1:200 dilution, R&D Systems, AF2018) was used as the indicator of SCs and rabbit anti-HA (1:200 dilution, Epizyme Biotech, LF315) was used as the indicator of HA expression. Alexa Fluor 488-conjugated donkey anti-mouse secondary IgG antibody (1:800 dilution, Invitrogen, A-21202), Alexa Fluor 555-conjugated donkey anti-rabbit secondary IgG antibody (1:800 dilution, Invitrogen, A-31570), and Alexa Fluor 647-conjugated donkey anti-goat secondary IgG antibody (1:800 dilution, Invitrogen, A-21447) were incubated in the dark for 2 h at room temperature after rinsing three times with PBS. The samples were then counterstained with DAPI (4,6-diamidino-2-phenylindole) as a nuclear indicator (Sigma, D9542). Images were acquired using a Leica confocal laser-scanning microscope (Leica, Wetzlar, Germany) with a 40× oil-immersion objective. All images were processed and analyzed using ImageJ software (National Institutes of Health, http://imagej.net/) and LAS X microcope software (Leica).

### Auditory testing

ABR measurements of mice were recorded on an RZ6 acoustic system (Tucker-Davis Technologies, Alachua, FL, USA) in a soundproof chamber^33^. Mice were anesthetized by intraperitoneal injection of 10 mg/kg xylazine and 100 mg/kg ketamine. ABR signals were recorded with three needle electrodes inserted into the mastoid portion (recording electrode), the subcutaneous tissue of the apex (reference electrode), and the rump (grounding electrode). ABR signals were elicited by 5 ms tone pips, amplified by 10,000, and pass-filtered with a 300 Hz–3 000 Hz passband. Mice were presented with tone burst stimuli of 5.6, 8, 11.3, 16, 22.6and 32 kHz at sound pressure levels between 20 dB and 90 dB in 5 dB increments until a threshold intensity that evoked a reproducible ABR waveform with an identifiable wave Ⅰ peak.

ABR measurements in cynomolgus monkeys was performed as previously described^28^. Briefly, cynomolgus monkeys were general anesthetized and anesthesia was maintained throughout the measurement. Electrodes were subdermally positioned at the central forehead and the ipsilateral mastoid, with the ground electrode placed at the contralateral mastoid. Closed-field ABRs to click and tone burst stimuli were recorded across the following frequencies: click, 0.5 kHz, 1 kHz, 2 kHz, 4 kHz, 8 kHz, and 16 kHz. Stimuli were presented at sound pressure levels ranging from 20 dB to 90 dB to determine hearing thresholds. Signals were processed using Neuro-Audio software, and body temperature was maintained with a heated pad within a stable range throughout the recording period. Following testing, anesthesia was discontinued. The animals were monitored closely until fully recovered from anesthesia and returned to its home cage.

### Anti-AAV NAb assay test

The anti-AAV neutralizing antibody (NAb) assay was conducted in HEK293T cells (ATCC, CRL-1573) for cynomolgus monkey serum samples. HEK293T cells were cultured in DMEM supplemented with 10% fetal bovine serum (FBS) and 1% penicillin/streptomycin and maintained in a 5% CO₂ atmosphere at 37 °C. 24 hours late, cells were seeded into 96-well plates at about 1.5 × 10⁴ cells per well and allowed to adhere for 4 hours. Heat-inactivated serum samples were serially diluted fivefold and combined with an equal volume of MAS012-CMV-Luc solution (Packgene Biotechnology), at a multiplicity of infection of 5 × 10¹⁰ vg/mL. The serum–virus mixtures were pre-incubated for 1 hour in a cell culture incubator before being added to the cell plates. Following 72 hours of incubation at 37 °C, the luciferase activity in the cell lysates was evaluated using a luciferase assay kit (Vazyme, DD1201-02) according to the manufacturer’s protocol. Luminescence, expressed as relative light units (RLU), was recorded on a Spark Multimode microplate reader (BioTek, Synergy2). The RLU values for test samples were compared against controls: a negative control (NC) containing medium without serum, and a blank control consisting of cells without MAS012-CMV-Luc. Percent inhibition was calculated using the following formula:

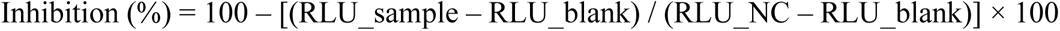

The NAb titer necessary for achieving 50% neutralization of vector transduction (ND₅₀) was calculated using nonlinear regression analysis in the Prism software (GraphPad Software Inc., San Diego, CA, USA).

### Integrin subunit coding gene analysis in cochlea

Single cell data was obtained from gEAR Portal (https://umgear.org/) in the datasets“scRNA-seq-P8, P12, P20 mouse cochleae^43^. Inspection of the data matrices confirmed that expression values were log1p-transformed and normalized. To visualize cell type–specific signatures, for scRNA-seq, gene expression was derived by computing the cell-number weighted average of stage-specific means (P8, P12, P20).

### Statistical analysis

Statistical analysis was performed with Prism 8 (GraphPad Software, La Jolla, CA, USA). Unpaired two-tailed Student’s t-tests were performed to compare differences between two groups, and comparison between multiple groups was performed by one-way analysis of variance (ANOVA) with Tukey’s post hoc test or Dunn’s multiple comparisons test. The exact replication numbers are stated in the figure legends. Data are presented as mean values ± s.e.m in the text; ns, not significant (*P* > 0.05); \**P* < 0.05, \*\**P* < 0.01, and \*\*\**P* < 0.001.

### Reporting summary

Further information on research design is available in the Nature Portfolio Reporting Summary linked to this article.

### Data availability

All data supporting the findings of this study are available within the paper and its Supplementary Information. Source data are provided with this paper.

## Acknowledgments

We thank Huixia Guo, Longlong Zhang, Zijing Wang, and Yuxin Zong for helpful discussions of the manuscript. We thank the Department of Laboratory Animal Resources Center of Fudan University for their support and assistance in animal husbandry. We thank Dr. Congcong Qi (Small Animal Behavior Monitoring Platform at the Laboratory Animal Center, Fudan University) for technical support in behavior tests. We thank the Core Facility of Shanghai Medical College, Fudan University, for their support and assistance in immunohistochemical experiments. This research was supported by National Key R&D Program of China (grant 2023YFA0915000 to B.Z. and H.T.), National Natural Science Foundation of China (grant 82225014, 82192864 to Y.S.; grant 82301318 to B.Z.; grant 82301332 to H.T.; grant 82501428 to CC), National Key R&D Program of China (grant 2021YFA1101302 to Y.S.), Science and Technology Commission of Shanghai Municipality (grant 23J31900100 to Y.S.), 2024 Annual Medical Service and Security Capacity Enhancement Project: National Key Clinical Specialty (to Y.S.), 2024 Healthcare Service and Assurance Capacity Enhancement Project: National Clinical Key Specialty (to Y.S.), New Cornerstone Science Foundation through the XPLORER PRIZE (to Y.S.), Shanghai Municipal Education Commission (grant 2023ZKZD12 to Y.S.), Shanghai Municipal Health Commission (grant 2024CXJQ02 to Y.S.), Science and Technology Innovation Program of Hunan Province (grant 2023RC4005 to Y.S.), Shanghai Science and Technology Program (grant 25S11900700 to H.W.), China Postdoctoral Science Foundation (grant 2023M740674 to Z.C.; grant 2025M771986 to C.C.), Science and Technology Commission of Shanghai (grant 24ZR1409500 to C.C.), Chenguang Program of Shanghai Education Development Foundation and Shanghai Municipal Education Commission (grant 23CGA08 to B.Z.), open research fund of Shanghai Key Laboratory of Gene Editing and Cell Therapy for Rare Diseases (grant gect-2025-Z14 to S.W.H; grant gect-2025-Z04 to C.C.).

## Author contributions

Y.S., and G.S., jointly supervised the project. Y.S., S.W.H and G.G. initiated the idea and conceived the study. S.W.H and G.G. designed and performed the in vitro studies. Y.Z., D.M., D.W., Z.C., B.Z., H.W., Q.S., H.Y., H.T., and Z.C. performed the in vitro studies. C.Y., Y.Z., Y.B., S.Z., C.C., X.F., S.H., L.H., X.H., X.W., Z.S., H.Y., and H.D. contributed to the in vivo study. S.W.H, C.Y., G.G., Y.Z., Y.B., S.Z., and C.C. wrote the paper, Y.S., G.S., S.W.H, C.Y., G.G., Y.Z., Y.B., S.Z., C.C., Z.C., B.Z., H.T., and H.L. reviewed and revised the manuscript. All authors read and approved the final manuscript.

## Extended Data

**Extended Data Fig. 1.**
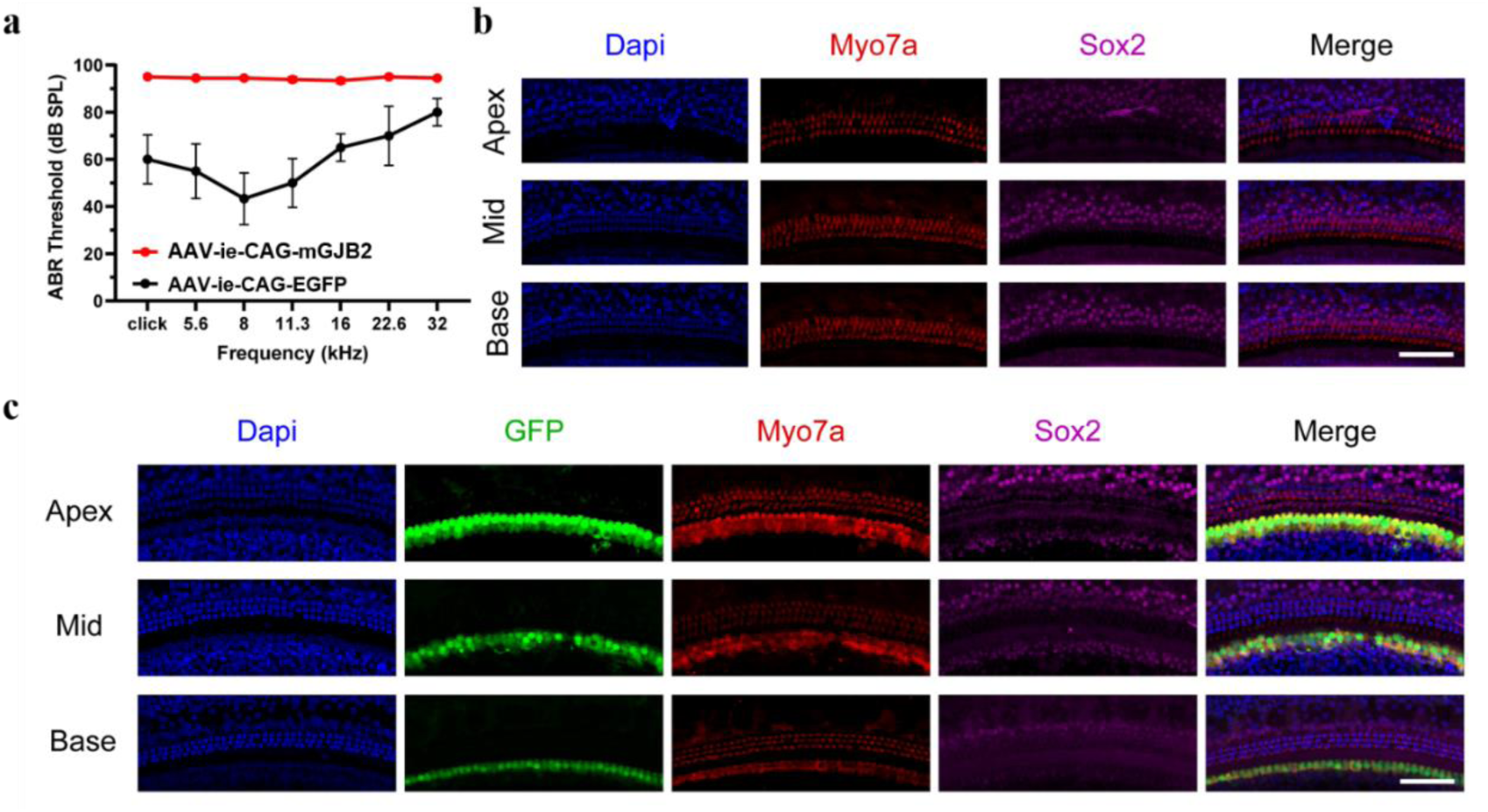
Ectopic expression of Cx26 in the inner ear leads to apoptosis of inner hair cells and subsequent hearing loss. **a**, Injection of AAV-ie-CAG-mGJB2 (titer: 1E13 vector genomes (VG)/mL) led to hearing loss in adult WT mice (n = 9), while administration of AAV-ie-CAG-EGFP (titer: 1E13 VG/mL) in the cochlea did not lead to hearing loss in adult WT mice (n = 3). **b**, Administration of AAV-ie-CAG-mGJB2 in the cochlea led to apoptosis of inner hair cells. **c**, Administration of AAV-ie-CAG-EGFP in the cochlea did not lead to cell death of inner hair cells. Blue: DAPI; Red: Myo7a; Green: GFP; Purple: Sox2. Scale bars: 100 μm.

**Extended Data Fig. 2.**
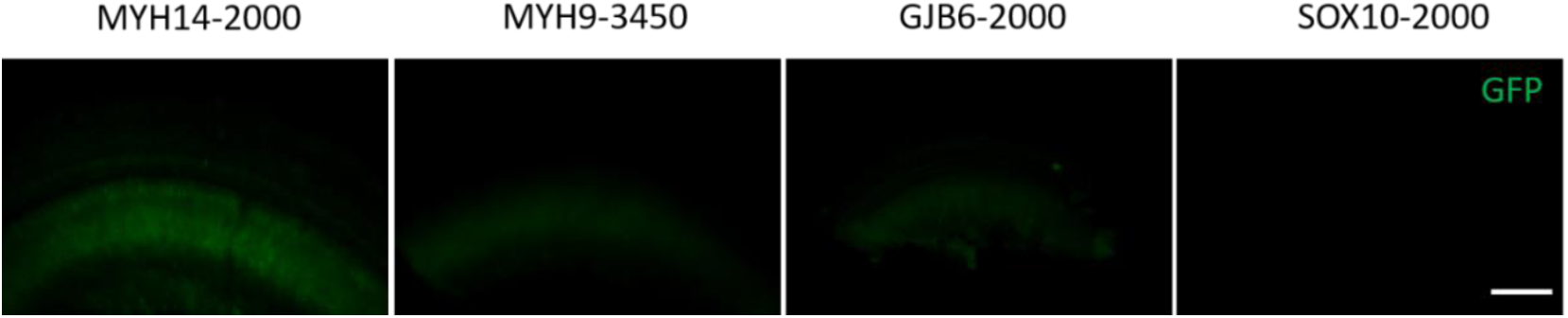
The efficiency of a subset of candidate cell type-specific promoters. The MYH14-2000 promoter drove the expression of GFP in the limbus spiralis in the inner ear, while the MYH9-3450, GJB6-2000, and SOX10-2000 promoters did not drive the expression of GFP in the inner ear. Green: GFP. Scale bars: 100 μm.

**Extended Data Fig. 3.**
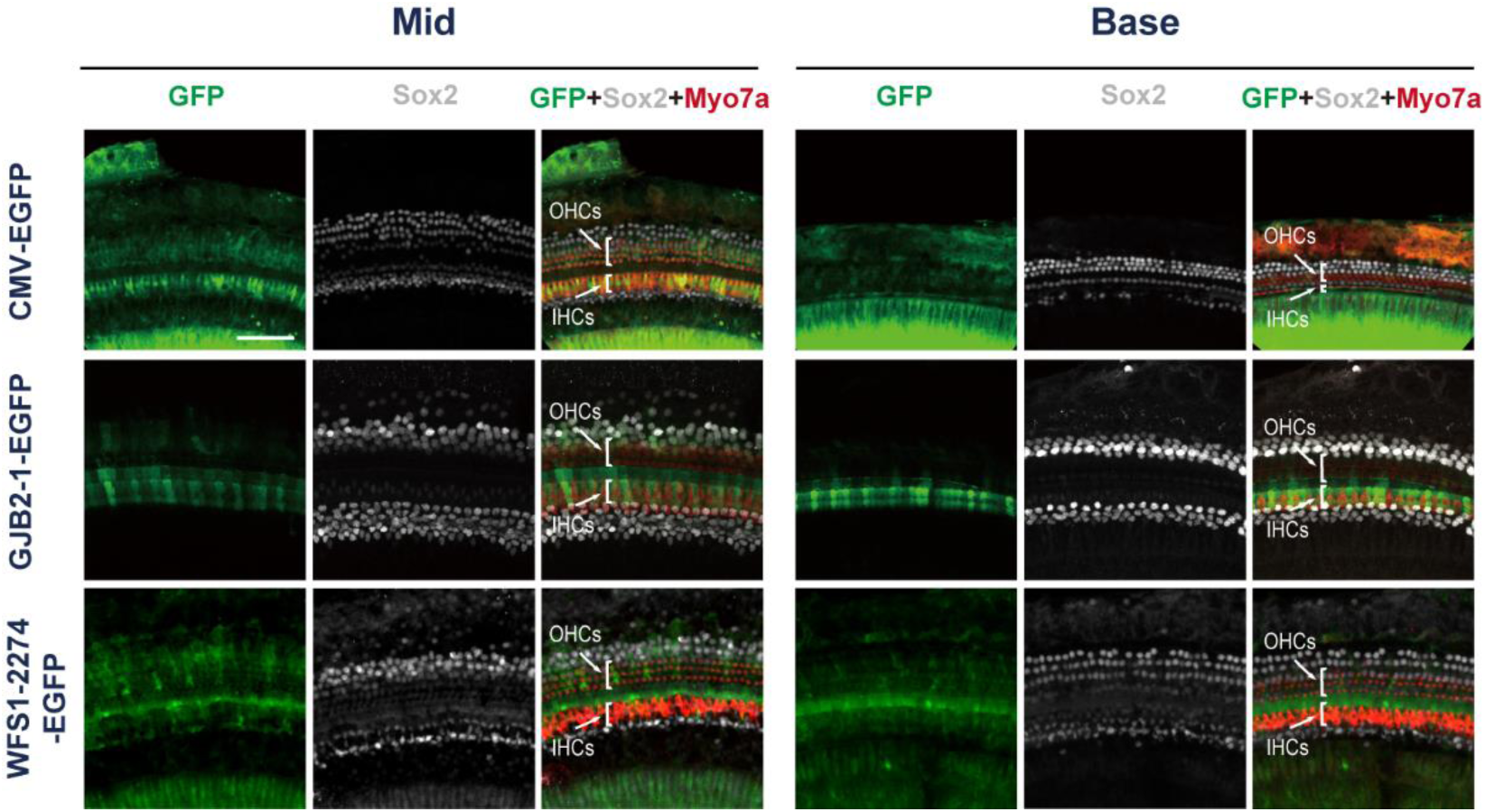
GJB2-1 promoter and WFS1-2274 promoter performed cell type-specific target in cochlear cells of newborn mice. The GJB2-1 promoter restricted expression of GFP to inner pillar cells (IPCs), outer pillar cells (OPCs), and Deiters’ cells (DCs) in the middle turn and base turn of the inner ear. In contrast, the WFS1-2274 promoter targeted a broader set of cell types, including Hensen’s cells (HeCs), DCs, OPCs, IPCs, and inner phalangeal cells (IPhCs) in the middle turn and base turn of the inner ear. Green: GFP; Gray: Sox2; Red: Myo7a. Scale bars: 100 μm.

**Extended Data Fig. 4.**
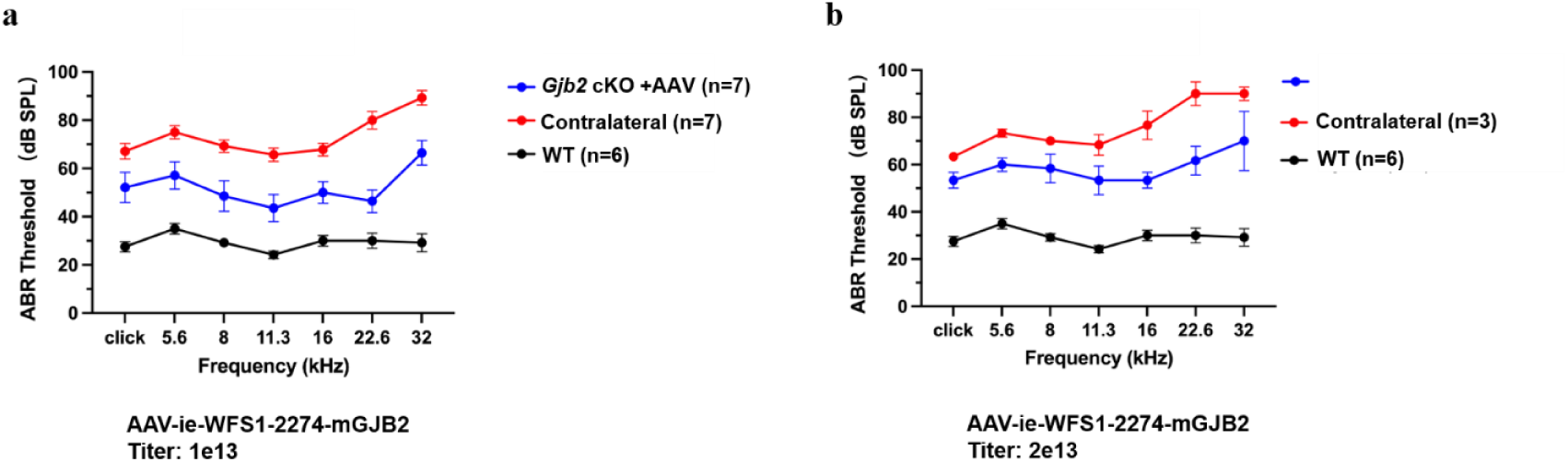
AAV-ie-WFS1-2274-mGJB2 partially restored hearing function in newborn Fgfr3-iCreERT2; Cx26^loxP/loxP^ mice. **a,** The ABR thresholds of newborn Fgfr3-iCreERT2; Cx26^loxP/loxP^ mice for click sound stimuli and pure-tone stimuli were recorded 4 weeks after injection of AAV-ie-GJB2-1-mGJB2 at P0–P2 (titer: 1E13 VG/mL) (WT, n = 6; injected Fgfr3-iCreERT2; Cx26^loxP/loxP^ mice, n = 7; contralateral ear, n = 7). **b,** The ABR thresholds of newborn Fgfr3-iCreERT2; Cx26^loxP/loxP^ mice for click sound stimuli and pure-tone stimuli were recorded 4 weeks after injection of AAV-ie-GJB2-1-mGJB2 at P0–P2 (titer: 2E13 VG/mL) (WT, n = 6; injected Fgfr3-iCreERT2; Cx26^loxP/loxP^ mice, n = 3; contralateral ear, n = 3).

**Extended Data Fig. 5.**
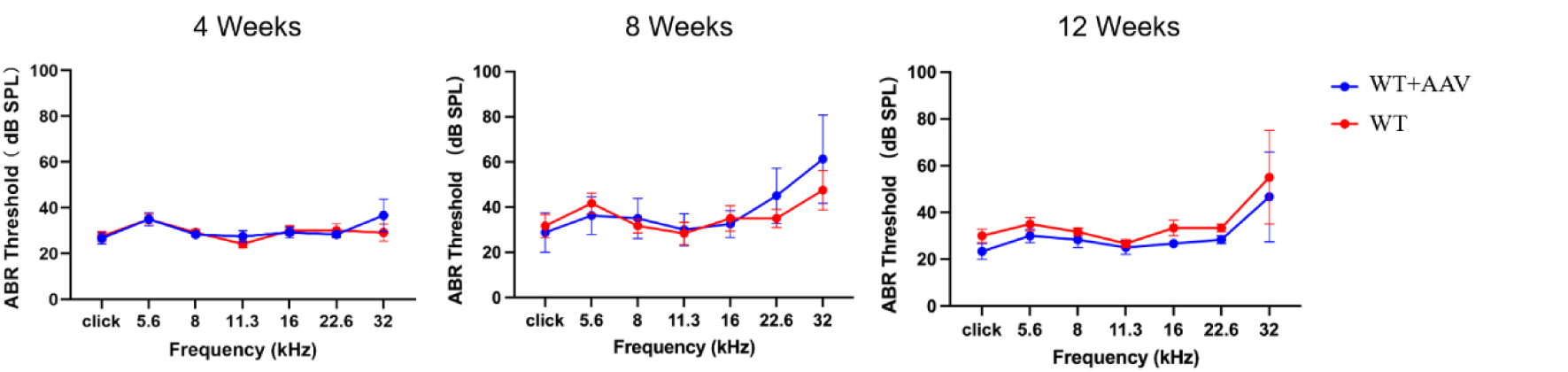
Evaluation of the safety of AAV-ie-GJB2-1-mGJB2 on hearing function. The ABR thresholds of WT mice for click sound stimuli and pure-tone stimuli were recorded 4 weeks (WT, n = 6; WT with AAV, n = 6), 8 weeks (WT, n = 6; WT with AAV, n = 6), and 12 weeks (WT, n = 3; WT with AAV, n = 3) after injection of AAV-ie-GJB2-1-mGJB2 at P0–P2.

**Extended Data Fig. 6.**
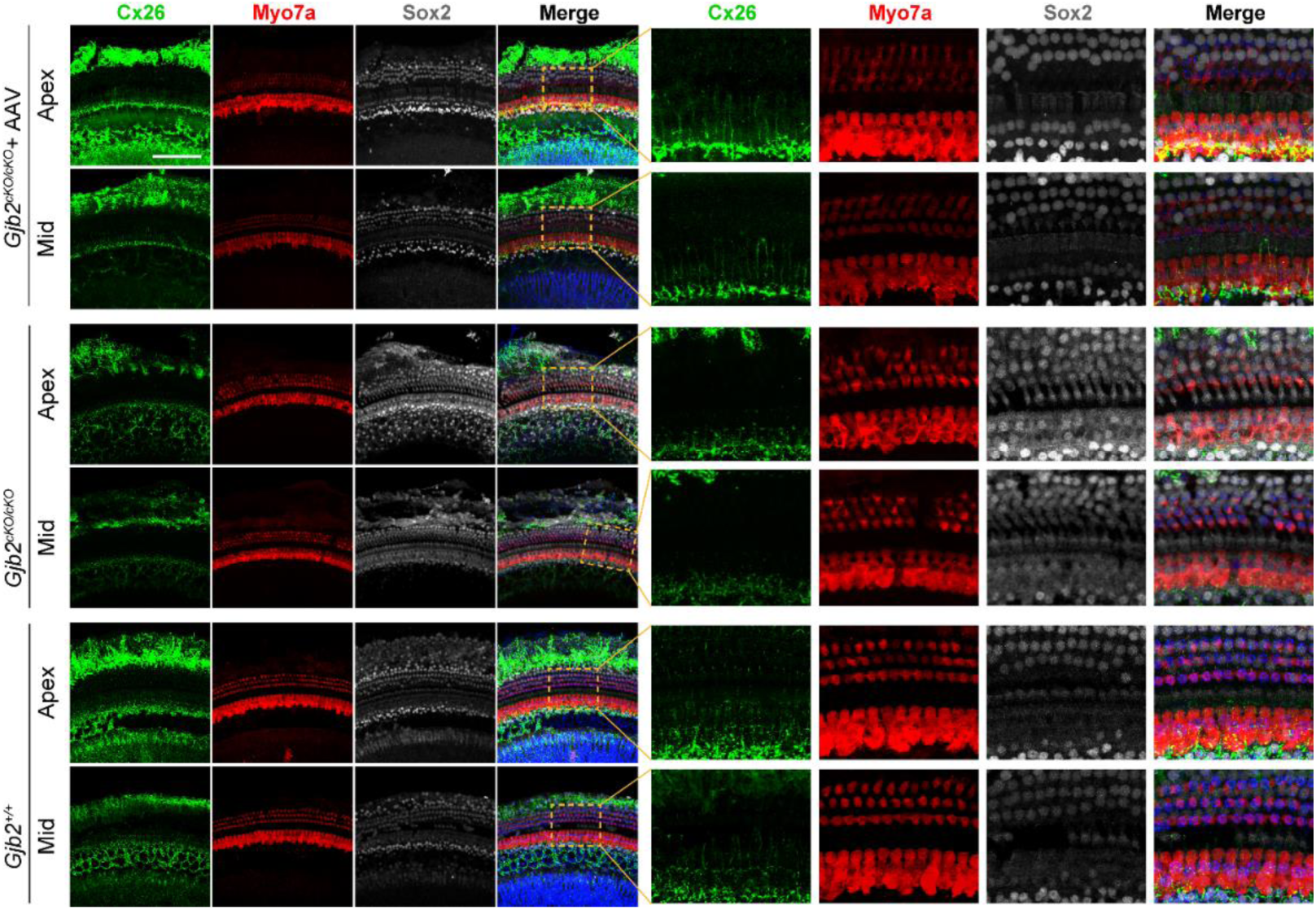
AAV-MAS012-GJB2-1-mGJB2 injection restored the expression of mouse Cx26 in apex and middle turns of the cochlea. The expression of Cx26 was decreased in PCs and DCs in the apex turn and middle turn in Fgfr3-iCreERT2; Cx26^loxP/loxP^ mice, and AAV-MAS012-GJB2-1-mGJB2 injection restored the expression of mouse Cx26 in PCs and DCs in the apex turn and middle turn in Fgfr3-iCreERT2; Cx26^loxP/loxP^ mice. The dashed boxes highlight the regions with restored expression of mouse Cx26 protein, and the right-hand panels show zoomed-in views of the areas within the dashed boxes. Green: Cx26; Red: Myo7a; Gray: Sox2; Blue: DAPI. Scale bars: 100 μm.

**Extended Data Fig. 7.**
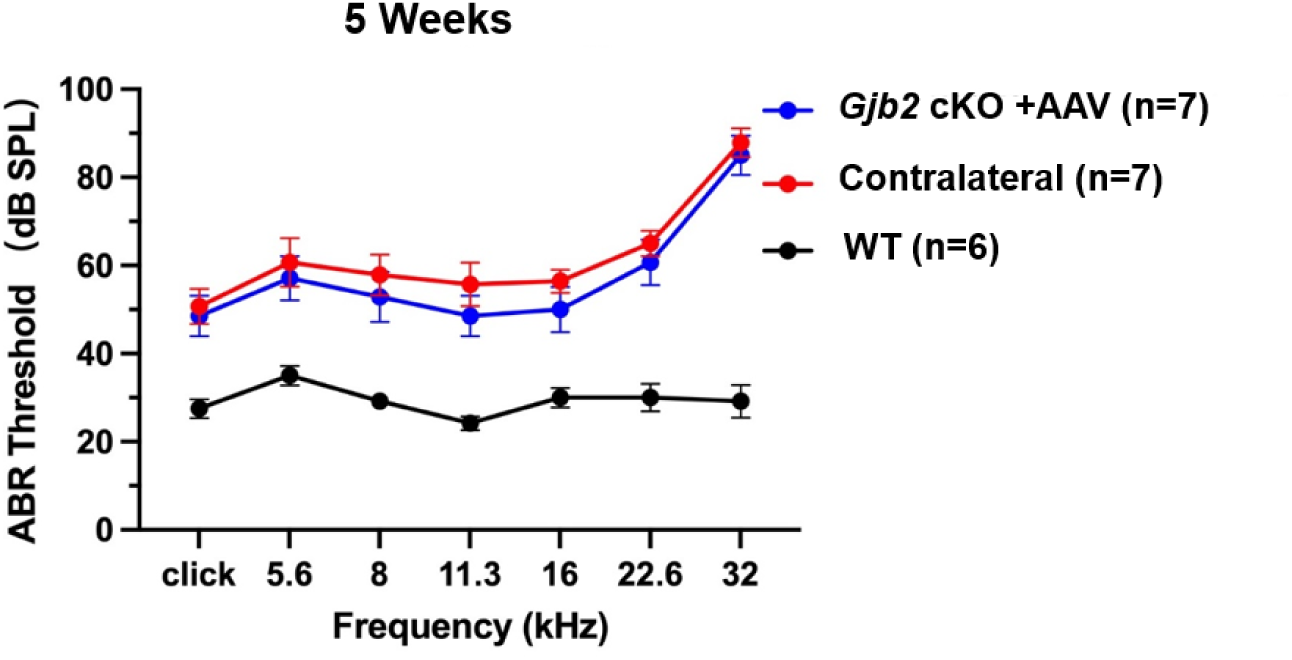
AAV-ie-GJB2-1-mGJB2 could not restore the hearing function of adult *Gjb2* cKO mice. Injection of AAV-ie-GJB2-1 pro-mGJB2 at P30 could not restore the hearing function of Gjb2 cKO mice at 5 weeks age (WT, n = 6; injected Gjb2 cKO mice, n = 7; contralateral ear, n = 7).

**Extended Data Fig. 8.**
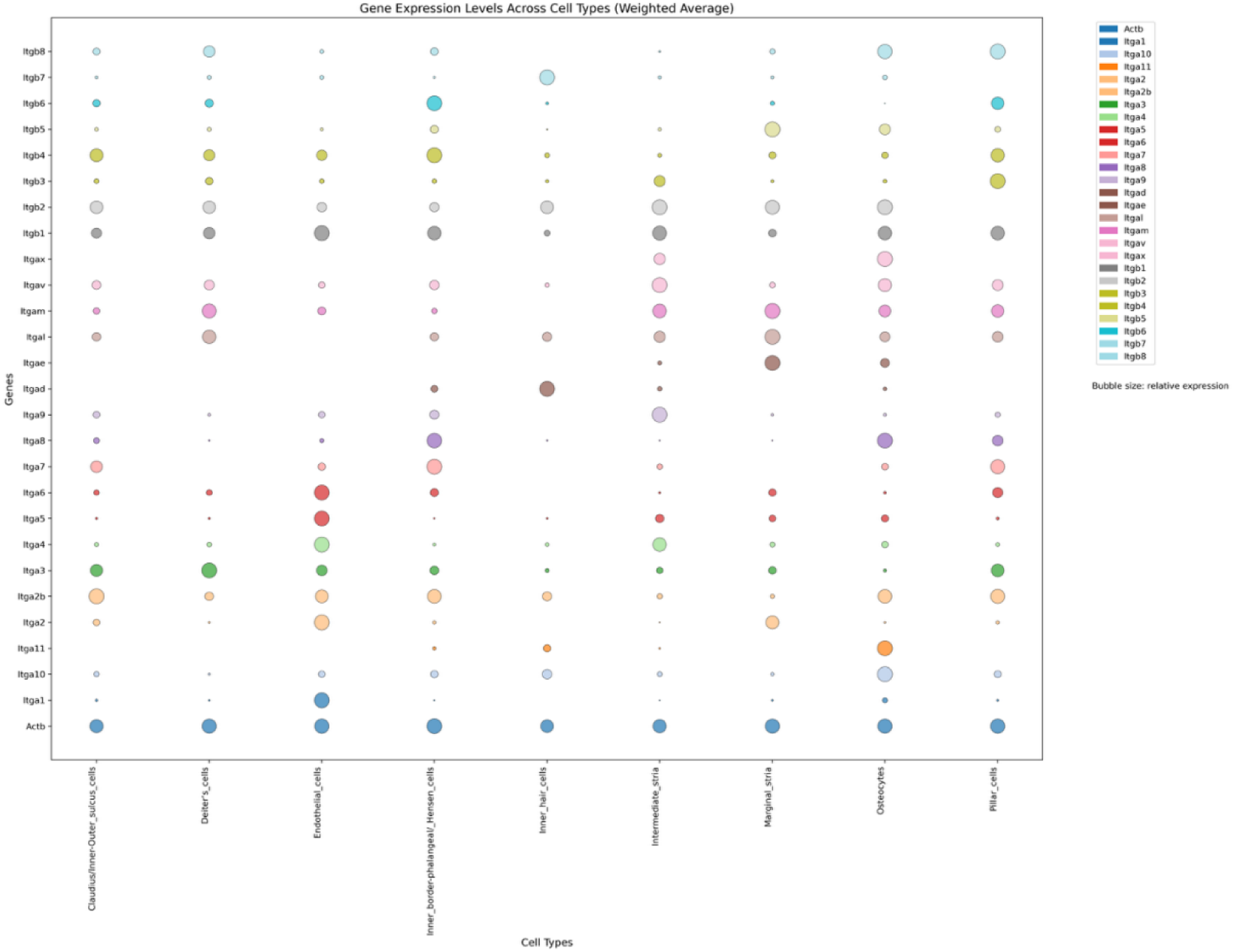
Bubble chart of integrin expression in different cell types in cochlea. Bubble chart depicting the relative expression levels integrin subunits in various cell types of the adult cochlea. The X-axis represents different cell types (Claudius/inner-outer-sulcus cells, Deiter’s cells, Endothelial cells, Inner border phalangeal cells/ Hensen cells, Intermediate stria, Marginal stria, and Pillar cells), while the Y-axis shows genes encoding integrin subunits. Expression levels were normalized to Actb as a housekeeping gene (arbitrary units) as bubble size. This analysis demonstrates that RGD integrin subunits are highly expressed across cochlear SCs and SVs, suggesting their potential involvement in gene delivery strategies.

**Extended Data Fig. 9.**
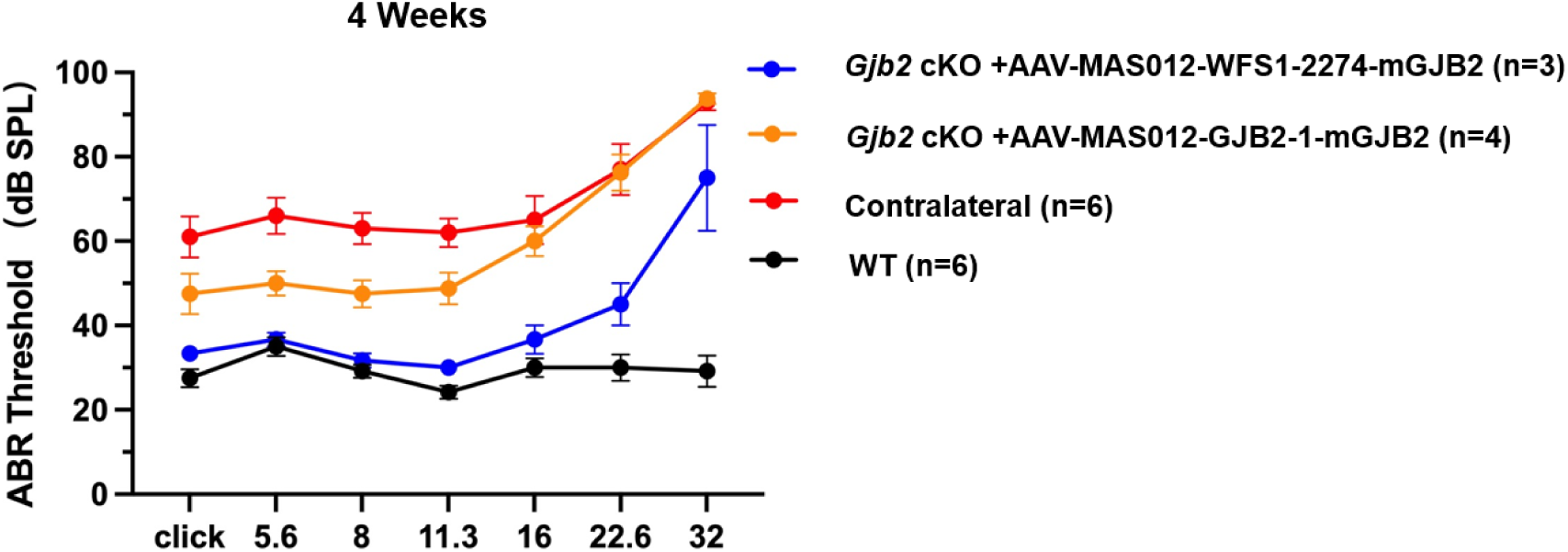
The AAV-MAS012-mediated GJB2 gene therapy systems restored the hearing function of P14 Fgfr3-iCreERT2; Cx26^loxP/loxP^ mice. The ABR thresholds of Fgfr3-iCreERT2; Cx26^loxP/loxP^ mice for click sound stimuli and pure-tone stimuli were recorded 2 weeks after injection of AAV-MAS012-GJB2-1-mGJB2 (titer: 1E13 VG/mL) and AAV-MAS012-WFS1-2274-mGJB2 (titer: 1E13 VG/mL) (WT, n = 6; AAV-MAS012-WFS1-2274-mGJB2–injected Gjb2 cKO mice, n = 3; AAV-MAS012-GJB2-1-mGJB2–injected Gjb2 cKO mice, n = 4; contralateral ear, n = 6) at P14.

**Extended Data Fig. 10.**
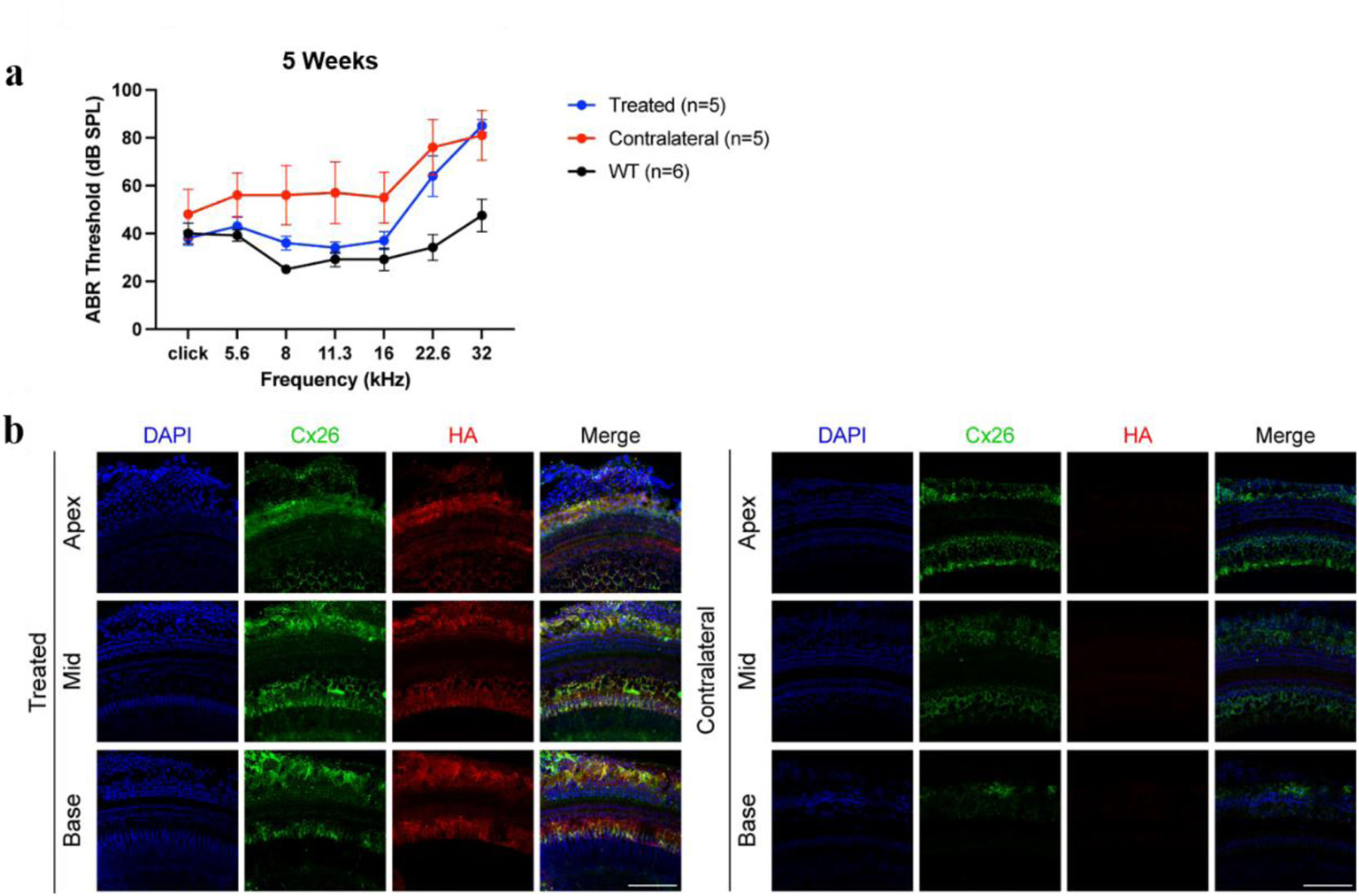
The exogenous Cx26 expression range was identified by adding the hemagglutinin (HA) tag to the C-terminus of human Cx26 in the AAV-mediated gene therapy system. **a,** Administration of AAV-MAS012-WFS1-2274-hGJB2-3×HA (titer: 3E13 VG/mL) at P30 rescued the hearing function of adult *Gjb2* cKO mice at 5 weeks of age (WT, n = 3; injected Fgfr3-iCreERT2; Cx26^loxP/loxP^ mouse ear, n = 4, contralateral ear, n = 4). **b,** The immunofluorescence results of the Cx26 showed the expression pattern of exogenous and endogenous Cx26 in the injected ear, while immunofluorescence results of the HA tag showed the expression pattern of exogenous Cx26 in the injected ear. Blue: DAPI; Green: Cx26; Red: HA-tag. Scale bars: 100 μm.

## References

1. Dror, A.A. & Avraham, K.B. Hearing Loss: Mechanisms Revealed by Genetics and Cell Biology. Annual Review of Genetics 43, 411–437 (2009).

2. Petit, C., Bonnet, C. & Safieddine, S. Deafness: from genetic architecture to gene therapy. Nat Rev Genet 24, 665–686 (2023).

3. Lieu, J.E.C., Kenna, M., Anne, S. & Davidson, L. Hearing Loss in Children: A Review. JAMA 324, 2195–2205 (2020).

4. Lv, J. et al. AAV1-hOTOF gene therapy for autosomal recessive deafness 9: a single-arm trial. Lancet (2024).

5. Wang, H. et al. Bilateral gene therapy in children with autosomal recessive deafness 9: single-arm trial results. Nature Medicine (2024).

6. Wang, H. et al. Hair cell-specific Myo15 promoter-mediated gene therapy rescues hearing in DFNB9 mouse model. Mol Ther Nucleic Acids 35, 102135 (2024).

7. Taiber, S., Gwilliam, K., Hertzano, R. & Avraham, K.B. The Genomics of Auditory Function and Disease. Annu Rev Genom Hum G 23, 275–299 (2022).

8. Shearer, A.E., Hildebrand, M.S., Odell, A.M. & Smith, R.J.H. in GeneReviews((R)). (eds. M.P. Adam et al.) (Seattle (WA); 1993).

9. Hu, S.W. et al. Engineering of the AAV-Compatible Hair Cell-Specific Small-Size Myo15 Promoter for Gene Therapy in the Inner Ear. Research-China 7 (2024).

10. Ivanchenko, M.V. & Corey, D.P. Finding a window for gene therapy for hereditary deafness. Proc Natl Acad Sci U S A 120, e2311864120 (2023).

11. Moisan, S., Le Nabec, A., Quillévéré, A., Le Maréchal, C. & Férec, C. Characterization of GJB2 cis-regulatory elements in the DFNB1 locus. Human Genetics 138, 1275–1286 (2019).

12. Guo, J. et al. GJB2 gene therapy and conditional deletion reveal developmental stage-dependent effects on inner ear structure and function. Mol Ther Methods Clin Dev 23, 319–333 (2021).

13. Takada, Y. et al. Connexin 26 null mice exhibit spiral ganglion degeneration that can be blocked by BDNF gene therapy. Hear Res 309, 124–135 (2014).

14. Yu, Q. et al. Virally expressed connexin26 restores gap junction function in the cochlea of conditional Gjb2 knockout mice. Gene Ther 21, 71–80 (2014).

15. Li, Q. et al. The pathogenesis of common Gjb2 mutations associated with human hereditary deafness in mice. Cell Mol Life Sci 80, 148 (2023).

16. Mehta, D. et al. Outcomes of evaluation and testing of 660 individuals with hearing loss in a pediatric genetics of hearing loss clinic. Am J Med Genet A 170, 2523–2530 (2016).

17. Carlyon, R.P. et al. Limitations on Temporal Processing by Cochlear Implant Users: A Compilation of Viewpoints. Trends Hear 29, 23312165251317006 (2025).

18. Zeng, F.G., Qi, J., Wu, C.C., Shu, Y. & Chai, R. Treating Hearing Loss: From Cochlear Implantation to Gene Therapy. Adv Sci (Weinh), e09960 (2025).

19. Cheng, X. et al. Gene Therapy vs Cochlear Implantation in Restoring Hearing Function and Speech Perception for Individuals With Congenital Deafness. JAMA Neurol (2025).

20. Wagner, H.J., Weber, W. & Fussenegger, M. Synthetic Biology: Emerging Concepts to Design and Advance Adeno-Associated Viral Vectors for Gene Therapy. Adv Sci (Weinh) 8, 2004018 (2021).

21. Tang, H.H. et al. Hearing of Otof-deficient mice restored by trans-splicing of N- and C-terminal otoferlin. Human Genetics (2022).

22. Qi, J. et al. AAV-Mediated Gene Therapy Restores Hearing in Patients with DFNB9 Deafness. Adv Sci (Weinh), e2306788 (2024).

23. Zhang, J. et al. Preliminary evidence for enhanced auditory cortex activation and mental development after gene therapy in children with autosomal recessive deafness 9. Nat Hum Behav 9, 1457–1469 (2025).

24. Valayannopoulos, V. et al. DB-OTO Gene Therapy for Inherited Deafness. N Engl J Med (2025).

25. Chen, S. et al. The spatial distribution pattern of Connexin26 expression in supporting cells and its role in outer hair cell survival. Cell Death Dis 9, 1180 (2018).

26. Kenneson, A., Van Naarden Braun, K. & Boyle, C. GJB2 (connexin 26) variants and nonsyndromic sensorineural hearing loss: a HuGE review. Genet Med 4, 258–274 (2002).

27. Chen, P. et al. Pathological mechanisms of connexin26-related hearing loss: Potassium recycling, ATP-calcium signaling, or energy supply? Front Mol Neurosci 15, 976388 (2022).

28. Sun, Q. et al. Combined AAV-mediated specific Gjb2 expression restores hearing in DFNB1 mouse models. Mol Ther 33, 3006–3021 (2025).

29. Ivanchenko, M.V. et al. Cell-specific delivery of GJB2 restores auditory function in mouse models of DFNB1 deafness and mediates appropriate expression in NHP cochlea. bioRxiv (2024).

30. Wang, X.H. et al. Viral-Mediated Connexin 26 Expression Combined with Dexamethasone Rescues Hearing in a Conditional Gjb2 Null Mice Model. Adv Sci (2024).

31. Seist, R., Copeland, J.S., Tao, L.T. & Groves, A.K. Rational design of a Lfng-enhancer AAV construct drives specific and efficient gene expression in inner ear supporting cells. Hearing Res 458 (2025).

32. Tan, F. et al. AAV-ie enables safe and efficient gene transfer to inner ear cells. Nat Commun 10, 3733 (2019).

33. Jiang, L. et al. Hearing restoration by gene replacement therapy for a multisite-expressed gene in a mouse model of human DFNB111 deafness. Am J Hum Genet 111, 2253–2264 (2024).

34. Hu, S.W. et al. PAM-flexible adenine base editing rescues hearing loss in a humanized MPZL2 mouse model harboring an East Asian founder mutation. Nat Commun 16, 7186 (2025).

35. Tu, Z.J. & Kiang, D.T. Mapping and characterization of the basal promoter of the human connexin26 gene. Biochim Biophys Acta 1443, 169–181 (1998).

36. Le Nabec, A. et al. 3D Chromatin Organization Involving MEIS1 Factor in the cis-Regulatory Landscape of GJB2. Int J Mol Sci 23 (2022).

37. Yoshimura, H., Nishio, S.Y. & Usami, S.I. Milestones toward cochlear gene therapy for patients with hereditary hearing loss. Laryngoscope Invest 6, 958–967 (2021).

38. Nishio, S.Y. et al. Gene expression profiles of the cochlea and vestibular endorgans: localization and function of genes causing deafness. Ann Otol Rhinol Laryngol 124 Suppl 1, 6S–48S (2015).

39. Molineris, I., Grassi, E., Ala, U., Di Cunto, F. & Provero, P. Evolution of Promoter Affinity for Transcription Factors in the Human Lineage. Mol Biol Evol 28, 2173–2183 (2011).

40. Lee, T.Y., Chang, W.C., Hsu, J.B.K., Chang, T.H. & Shien, D.M. GPMiner: an integrated system for mining combinatorial cis-regulatory elements in mammalian gene group. Bmc Genomics 13 (2012).

41. Wang, X.H., et al. PARP inhibitor rescues hearing and hair cell impairment in Cx26-null mice. View-China 4 (2023).

42. Tao, Y. et al. AAV-ie-K558R mediated cochlear gene therapy and hair cell regeneration. Signal Transduct Tar 7 (2022).

43. Jean, P. et al. Single-cell transcriptomic profiling of the mouse cochlea: An atlas for targeted therapies. Proc Natl Acad Sci U S A 120, e2221744120 (2023).

44. Tabebordbar, M. et al. Directed evolution of a family of AAV capsid variants enabling potent muscle-directed gene delivery across species. Cell 184, 4919–4938 e4922 (2021).

45. Cui, C., et al. A base editor for the long-term restoration of auditory function in mice with recessive profound deafness. Nature Biomedical Engineering 9 (2025).

46. Zhang, L. et al. Virally mediated connexin 26 expression in postnatal scala media significantly and transiently preserves hearing in connexin 30 null mice (vol 10, 900416, 2022). Frontiers in Cell and Developmental Biology 10 (2022).

47. Sun, Y. et al. Connexin30 null and conditional connexin26 null mice display distinct pattern and time course of cellular degeneration in the cochlea. J Comp Neurol 516, 569–579 (2009).

48. Ahmad, S. et al. Restoration of connexin26 protein level in the cochlea completely rescues hearing in a mouse model of human connexin30-linked deafness. Proc Natl Acad Sci U S A 104, 1337–1341 (2007).

49. Teubner, B. et al. Connexin30 (Gjb6)-deficiency causes severe hearing impairment and lack of endocochlear potential. Hum Mol Genet 12, 13–21 (2003).

50. Zhang, L. et al. Preclinical evaluation of the efficacy and safety of AAV1-hOTOF in mice and nonhuman primates. Mol Ther Methods Clin Dev 31, 101154 (2023).

51. Nettelbeck, D.M., Jerome, V. & Muller, R. A strategy for enhancing the transcriptional activity of weak cell type-specific promoters. Gene Ther 5, 1656–1664 (1998).

52. Mason, J.B. et al. Delivery and evaluation of recombinant adeno-associated viral vectors in the equine distal extremity for the treatment of laminitis. Equine Vet J 49, 79–86 (2017).

53. Fang, K.L. et al. A comprehensive study of AAV tropism across C57BL/6 mice, BALB/c mice, and crab-eating macaques. Mol Ther Meth Clin D 33 (2025).

