## Extended Data for "Combination of engineered cell type-specific promoters and a high-efficiency AAV capsid restores hearing in adult DFNB1 mice model with demonstrated safety in nonhuman primate"

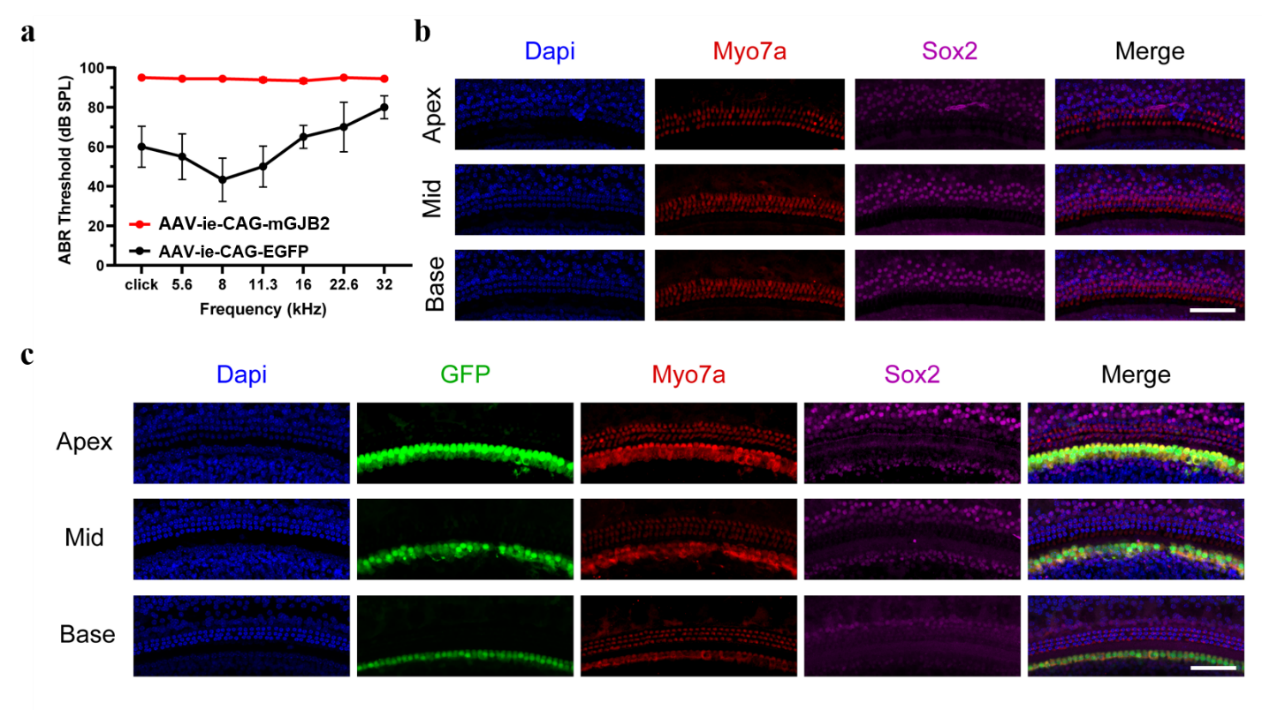

**Extended Data Fig. 1** **Ectopic expression of Cx26 in the inner ear leads to apoptosis of inner hair cells and subsequent hearing loss.** **a**, Injection of AAV-ie-CAG-mGJB2 (titer: 1E13 vector genomes (VG)/mL) led to hearing loss in adult WT mice (n = 9), while administration of AAV-ie-CAG-EGFP (titer: 1E13 VG/mL) in the cochlea did not lead to hearing loss in adult WT mice (n = 3). **b**, Administration of AAV-ie-CAG-mGJB2 in the cochlea led to apoptosis of inner hair cells. **c**, Administration of AAV-ie-CAG-EGFP in the cochlea did not lead to cell death of inner hair cells. Blue: DAPI; Red: Myo7a; Green: GFP; Purple: Sox2. Scale bars: 100 μm.

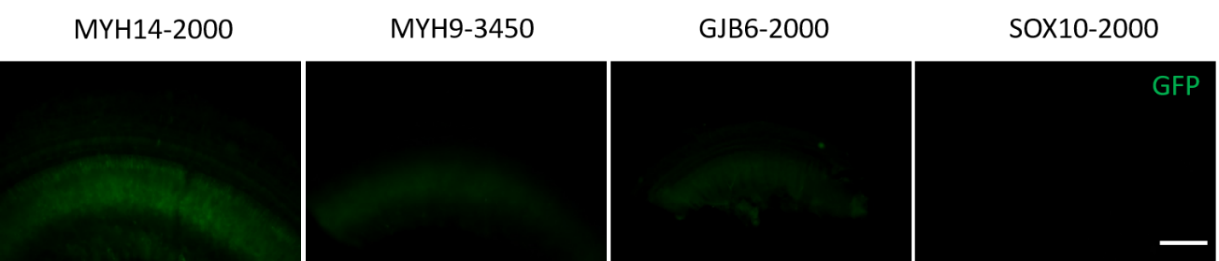

**Extended Data Fig. 2** **The efficiency of a subset of candidate cell type-specific promoters.** The MYH14-2000 promoter drove the expression of GFP in the limbus spiralis in the inner ear, while the MYH9-3450, GJB6-2000, and SOX10-2000 promoters did not drive the expression of GFP in the inner ear. Green: GFP. Scale bars: 100 μm.

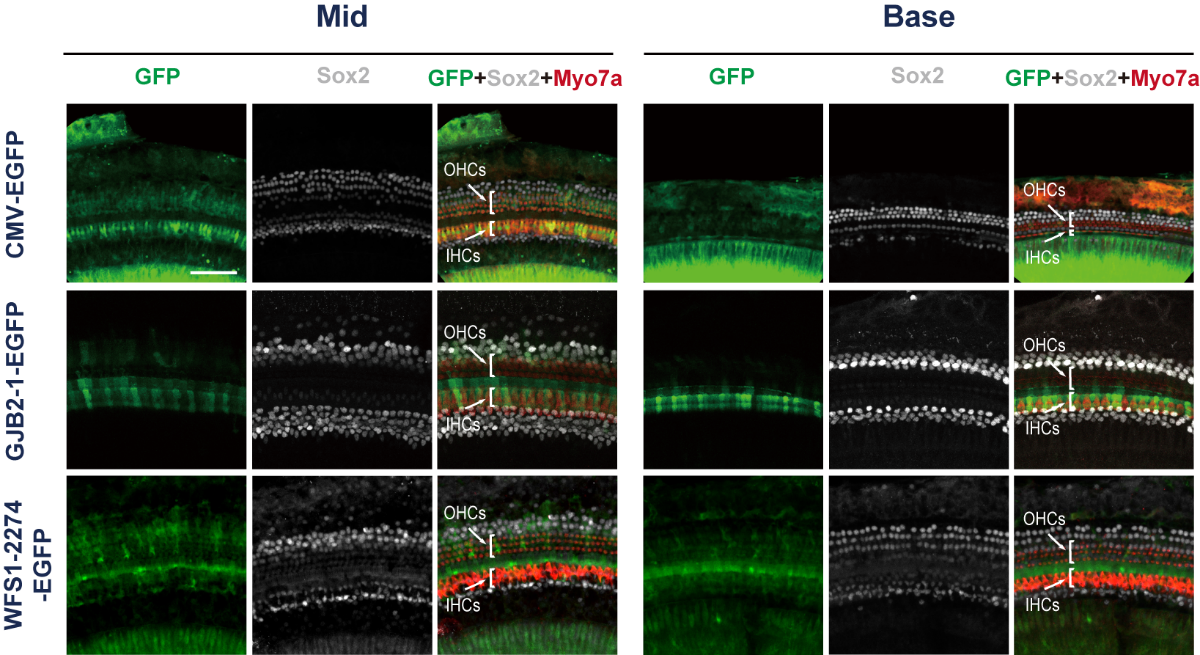

**Extended Data Fig. 3 GJB2-1 promoter and WFS1-2274 promoter performed cell type-specific target in cochlear cells of newborn mice.** The GJB2-1 promoter restricted expression of GFP to inner pillar cells (IPCs), outer pillar cells (OPCs), and Deiters’ cells (DCs) in the middle turn and base turn of the inner ear. In contrast, the WFS1-2274 promoter targeted a broader set of cell types, including Hensen's cells (HeCs), DCs, OPCs, IPCs, and inner phalangeal cells (IPhCs) in the middle turn and base turn of the inner ear. Green: GFP; Gray: Sox2; Red: Myo7a. Scale bars: 100 μm.

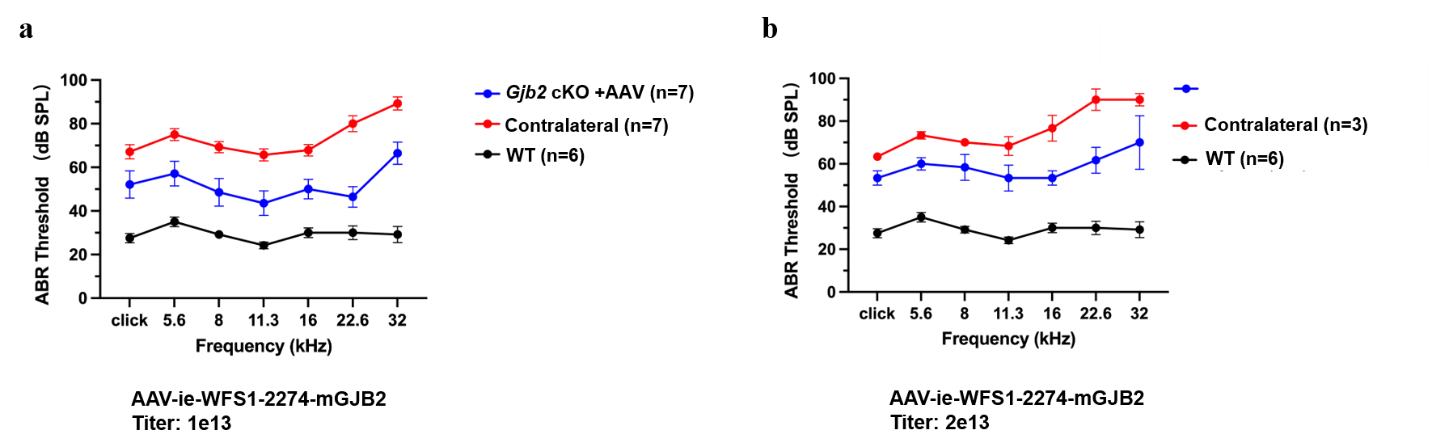

**Extended Data Fig. 4 AAV-ie-WFS1-2274-mGJB2 partially restored hearing function in newborn Fgfr3-iCreERT2; Cx26^loxP/loxP^ mice. a,** The ABR thresholds of newborn Fgfr3-iCreERT2; Cx26^loxP/loxP^ mice for click sound stimuli and pure-tone stimuli were recorded 4 weeks after injection of AAV-ie-GJB2-1-mGJB2 at P0–P2 (titer: 1E13 VG/mL) (WT, n = 6; injected Fgfr3-iCreERT2; Cx26^loxP/loxP^ mice, n = 7; contralateral ear, n = 7). **b,** The ABR thresholds of newborn Fgfr3-iCreERT2; Cx26^loxP/loxP^ mice for click sound stimuli and pure-tone stimuli were recorded 4 weeks after injection of AAV-ie-GJB2-1-mGJB2 at P0–P2 (titer: 2E13 VG/mL) (WT, n = 6; injected Fgfr3-iCreERT2; Cx26^loxP/loxP^ mice, n = 3; contralateral ear, n = 3).

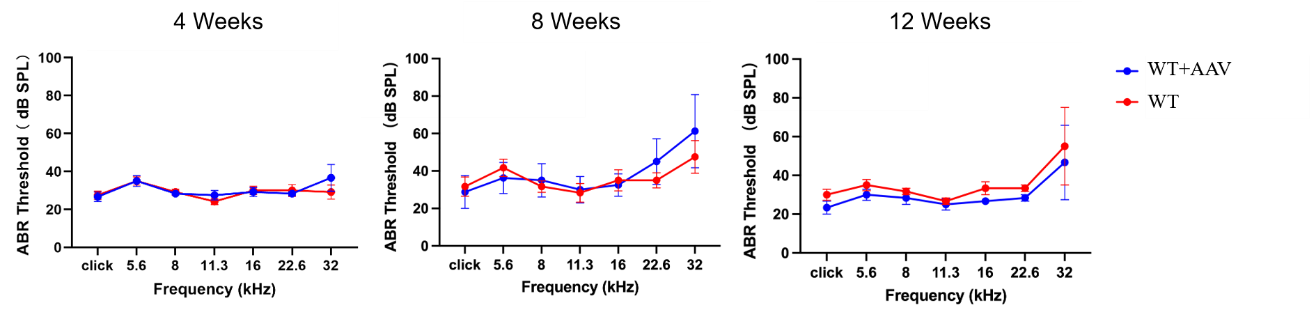

**Extended Data Fig. 5 Evaluation of the safety of AAV-ie-GJB2-1-mGJB2 on hearing function.** The ABR thresholds of WT mice for click sound stimuli and pure-tone stimuli were recorded 4 weeks (WT, n = 6; WT with AAV, n = 6), 8 weeks (WT, n = 6; WT with AAV, n = 6), and 12 weeks (WT, n = 3; WT with AAV, n = 3) after injection of AAV-ie-GJB2-1-mGJB2 at P0–P2.

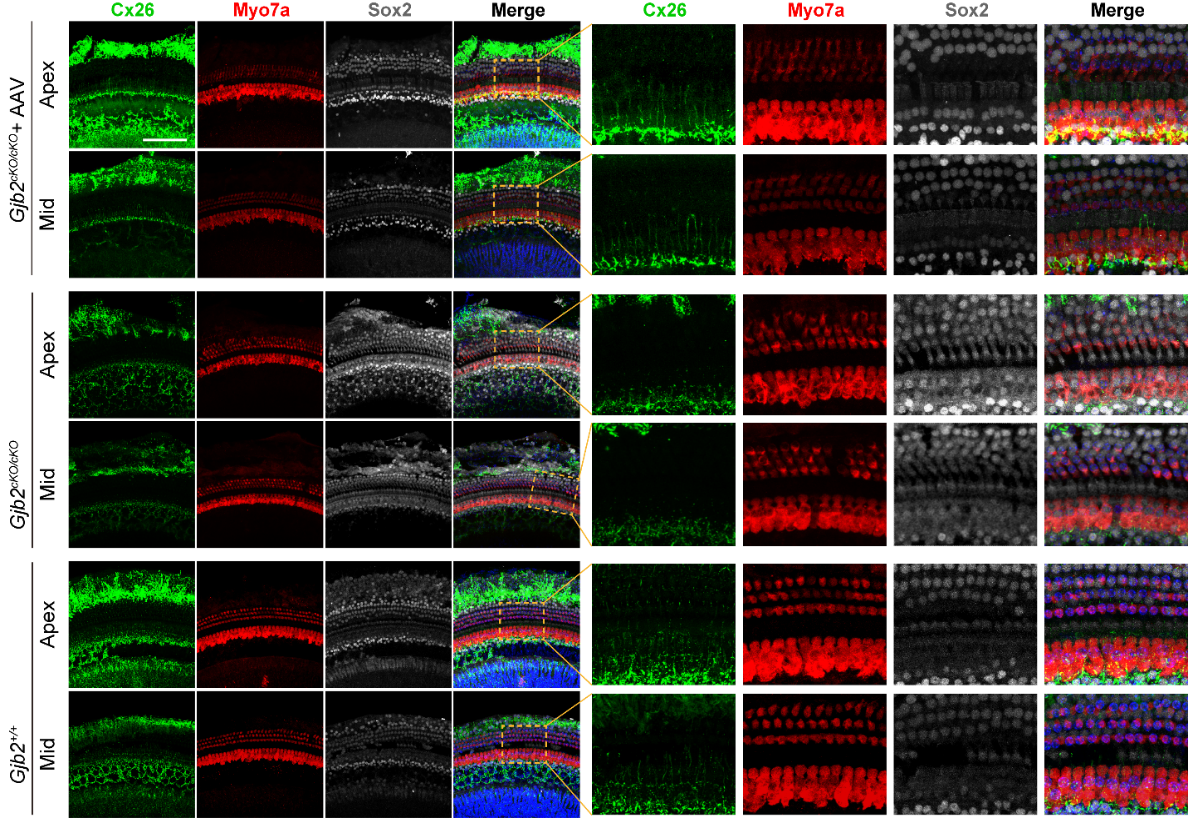

**Extended Data Fig. 6 AAV-MAS012-GJB2-1-mGJB2 injection restored the expression of mouse Cx26 in apex and middle turns of the cochlea.** The expression of Cx26 was decreased in PCs and DCs in the apex turn and middle turn in Fgfr3-iCreERT2; Cx26^loxP/loxP^ mice, and AAV-MAS012-GJB2-1-mGJB2 injection restored the expression of mouse Cx26 in PCs and DCs in the apex turn and middle turn in Fgfr3-iCreERT2; Cx26^loxP/loxP^ mice. The dashed boxes highlight the regions with restored expression of mouse Cx26 protein, and the right-hand panels show zoomed-in views of the areas within the dashed boxes. Green: Cx26; Red: Myo7a; Gray: Sox2; Blue: DAPI. Scale bars: 100 μm.

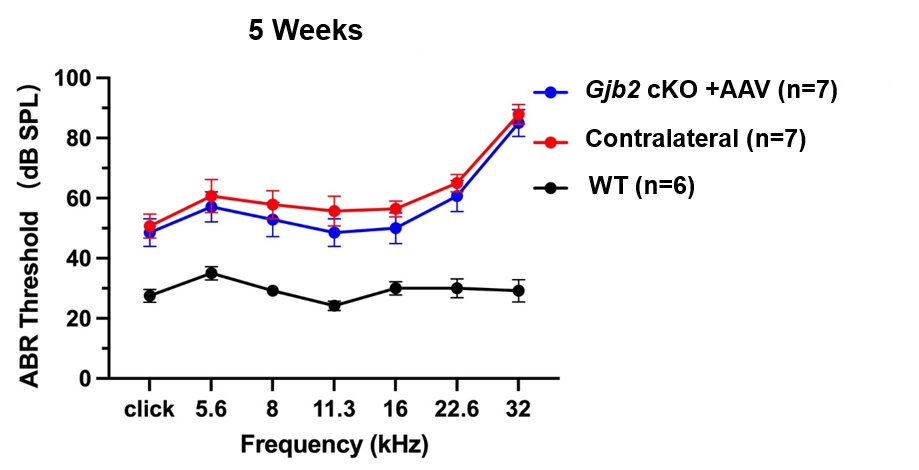

**Extended Data Fig. 7 AAV-ie-GJB2-1-mGJB2 could not restore the hearing function of adult *Gjb2* cKO mice.** Injection of AAV-ie-GJB2-1 pro-mGJB2 at P30 could not restore the hearing function of Gjb2 cKO mice at 5 weeks age (WT, n = 6; injected Gjb2 cKO mice, n = 7; contralateral ear, n = 7).

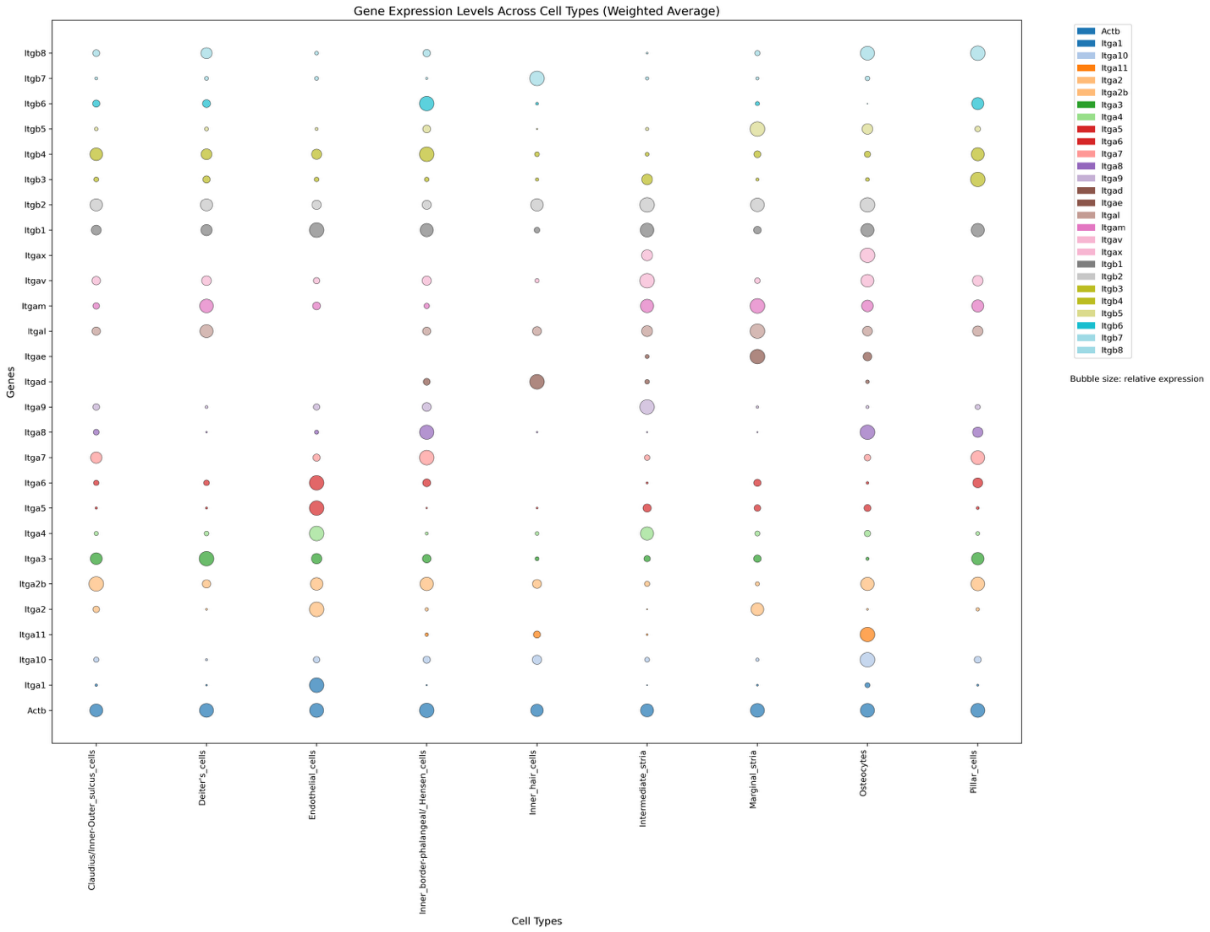

**Extended Data Fig. 8** **Bubble chart of integrin expression in different cell types in cochlea.** Bubble chart depicting the relative expression levels integrin subunits in various cell types of the adult cochlea. The X-axis represents different cell types (Claudius/inner-outer-sulcus cells, Deiter’s cells, Endothelial cells, Inner border phalangeal cells/ Hensen cells, Intermediate stria, Marginal stria, and Pillar cells), while the Y-axis shows genes encoding integrin subunits. Expression levels were normalized to Actb as a housekeeping gene (arbitrary units) as bubble size. This analysis demonstrates that RGD integrin subunits are highly expressed across cochlear SCs and SVs, suggesting their potential involvement in gene delivery strategies.

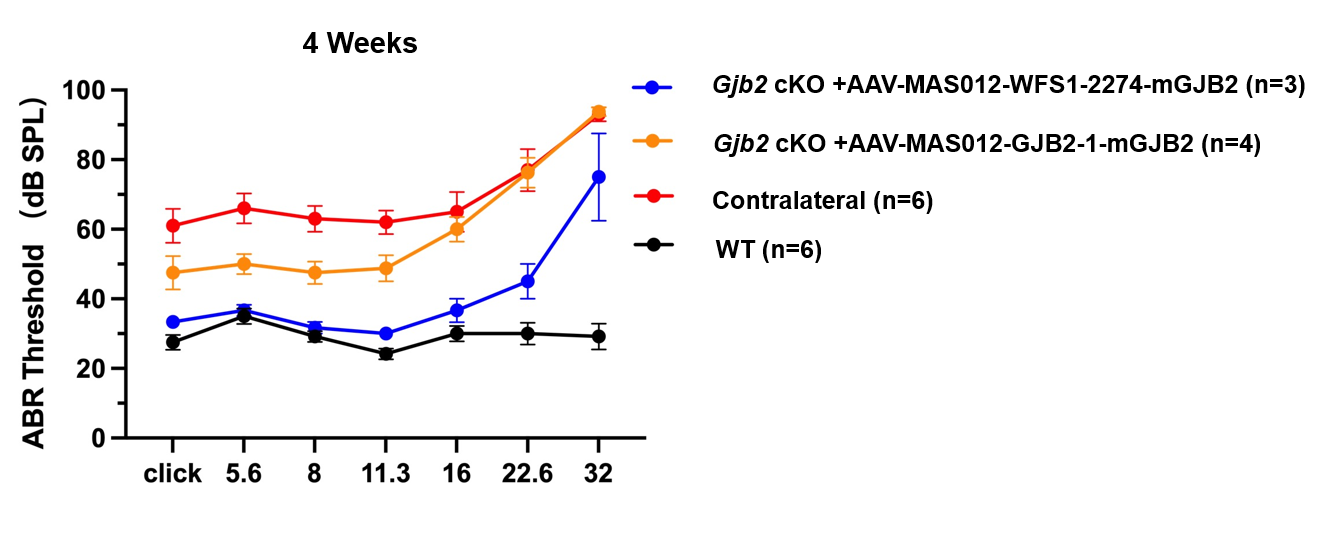

**Extended Data Fig. 9 The AAV-MAS012-mediated GJB2 gene therapy systems restored the hearing function of P14 Fgfr3-iCreERT2; Cx26****^loxP/loxP^ mice.** The ABR thresholds of Fgfr3-iCreERT2; Cx26^loxP/loxP^ mice for click sound stimuli and pure-tone stimuli were recorded 2 weeks after injection of AAV-MAS012-GJB2-1-mGJB2 (titer: 1E13 VG/mL) and AAV-MAS012-WFS1-2274-mGJB2 (titer: 1E13 VG/mL) (WT, n = 6; AAV-MAS012-WFS1-2274-mGJB2–injected Gjb2 cKO mice, n = 3; AAV-MAS012-GJB2-1-mGJB2–injected Gjb2 cKO mice, n = 4; contralateral ear, n = 6) at P14.

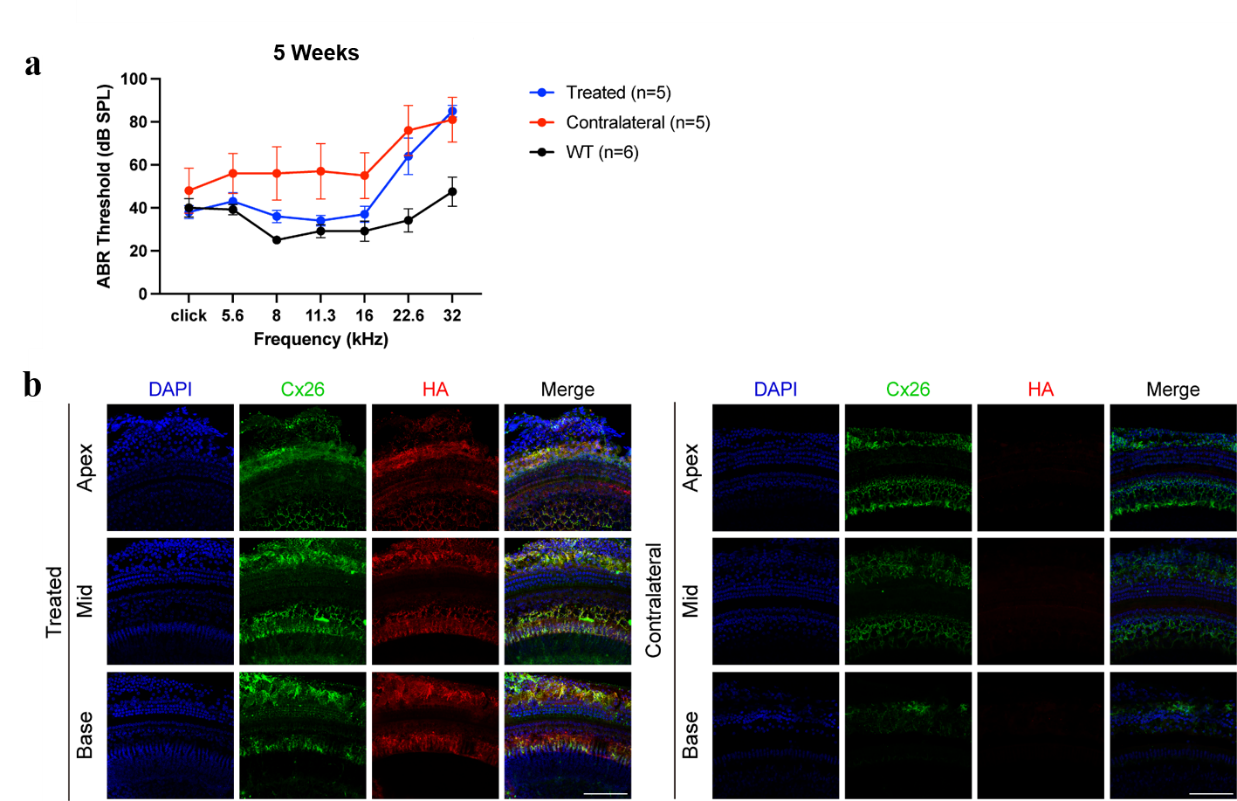

**Extended Data Fig. 10 The exogenous Cx26 expression range was identified by adding the hemagglutinin (HA) tag to the C-terminus of human Cx26 in the AAV-mediated gene therapy system. a,** Administration of AAV-MAS012-WFS1-2274-hGJB2-3×HA (titer: 3E13 VG/mL) at P30 rescued the hearing function of adult *Gjb2* cKO mice at 5 weeks of age (WT, n = 3; injected Fgfr3-iCreERT2; Cx26^loxP/loxP^ mouse ear, n = 4, contralateral ear, n = 4). **b,** The immunofluorescence results of the Cx26 showed the expression pattern of exogenous and endogenous Cx26 in the injected ear, while immunofluorescence results of the HA tag showed the expression pattern of exogenous Cx26 in the injected ear. Blue: DAPI; Green: Cx26; Red: HA-tag. Scale bars: 100 μm.
