## Supplementary Information for "Combination of engineered cell type-specific promoters and a high-efficiency AAV capsid restores hearing in adult DFNB1 mice model with demonstrated safety in nonhuman primate"

**Supplementary Table 1. Hereditary deafness genes with similar expression ranges to *Gjb2*.**

|  | IDC | SL | ISC | SV | SLig | SP | ESC | CC | HeC | SC | PC | IHC | OHC | TM | RM |
| --- | --- | --- | --- | --- | --- | --- | --- | --- | --- | --- | --- | --- | --- | --- | --- |
| *Gjb2* | √ | √ | √ | √ | √ | √ | √ | √ | √ | √ | √ | 🞨 | 🞨 | 🞨 | 🞨 |
| *Gjb6* | √ | √ | √ | √ | √ | √ | √ | √ | √ | √ | √ | 🞨 | 🞨 | 🞨 | 🞨 |
| *Myh14* | 🞨 | 🞨 | 🞨 | √ | √ | √ | √ | √ | √ | √ | √ | √ | √ | 🞨 | 🞨 |
| *Wfs1* | √ | √ | √ | √ | √ | 🞨 | √ | √ | √ | √ | √ | √ | √ | 🞨 | √ |
| *Eya1* | √ | √ | √ | √ | √ | 🞨 | 🞨 | √ | √ | √ | √ | √ | √ | 🞨 | √ |
| *Myh9* | 🞨 | √ | √ | 🞨 | √ | √ | √ | √ | √ | √ | √ | 🞨 | √ | 🞨 | √ |
| *Sox10* | √ | 🞨 | √ | √ | 🞨 | √ | √ | √ | √ | √ | √ | 🞨 | 🞨 | 🞨 | √ |
| *Eya4* | 🞨 | √ | √ | 🞨 | 🞨 | √ | √ | √ | √ | 🞨 | 🞨 | 🞨 | 🞨 | 🞨 | 🞨 |
| *Time* | 🞨 | 🞨 | 🞨 | √ | 🞨 | 🞨 | 🞨 | √ | √ | √ | √ | √ | √ | 🞨 | √ |
| *Adcy1* | 🞨 | 🞨 | √ | 🞨 | 🞨 | 🞨 | 🞨 | √ | √ | √ | 🞨 | √ | √ | 🞨 | 🞨 |
| *Serpinb6* | 🞨 | √ | √ | √ | √ | 🞨 | √ | √ | 🞨 | 🞨 | 🞨 | √ | √ | 🞨 | 🞨 |
| *Col4a3* | 🞨 | √ | √ | √ | √ | √ | √ | 🞨 | 🞨 | 🞨 | 🞨 | 🞨 | 🞨 | 🞨 | √ |
| *Col4a5* | 🞨 | √ | √ | √ | √ | √ | √ | 🞨 | 🞨 | 🞨 | 🞨 | 🞨 | 🞨 | 🞨 | √ |
| *Cdh7* | √ | √ | 🞨 | √ | √ | √ | 🞨 | 🞨 | 🞨 | 🞨 | √ | √ | √ | 🞨 | 🞨 |
| *Cldn14* | 🞨 | 🞨 | 🞨 | 🞨 | 🞨 | 🞨 | 🞨 | √ | √ | √ | √ | √ | √ | √ | √ |

Abbreviations: IDC, inter dental cell; SL, spiral limbus; ISC, inner sulcus cell; SV, stria vascularis; SLig, Spiral ligament; SP, spiral prominence; ESC, external sulcus cell; CC, Claudius’ cell; HeC, Hensen’s cell; SC, supporting cell; PC, pillar cell; IHC, inner hair cell; OHC, outer hair cell; TM, Tectorial membrane; RM, Reissner’s membrane.

**Supplementary Table 2. Target range of the 21 cell type-specific promoter candidates.**

| Promoters | Expression of reporter gene | Range |
| --- | --- | --- |
| GJB2-1 | Yes | IPC OPC DC |
| GJB2-2 | No | / |
| GJB2-Y1 | No | / |
| GJB2-Y2 | No | / |
| GJB2-Y3 | No | / |
| GJB2-Y4 | No | / |
| GJB2-Y5 | No | / |
| hGJB2-2 | No | / |
| hGJB2-10 | No | / |
| GJB6-2000 | No | / |
| GJB6-3450 | Yes | SL |
| MYH14-2000 | Yes | SL |
| MYH14-3450 | No | / |
| WFS1-2274 | Yes | IBC IPhC IPC OPC DC HeC SV |
| WFS1-3450 | No | / |
| EYA1-2000 | No | / |
| EYA1-3450 | No | / |
| MYH9-2000 | No | / |
| MYH9-3450 | Yes | SL |
| SOX10-2000 | No | / |
| SOX10-3450 | No | / |

Abbreviations: IPC, inner pillar cell; OPC, outer pillar cell; DC, Deiter's cell; IBC, inner border cell; IPhC, inner phalangeal cell; HeC, Hensen's cell; SV, stria vascularis; SL, spiral limbus.

**Supplementary Table 3. Parameters of** **hematology, coagulation, and comprehensive serum biochemistry in cynomolgus monkeys at multiple time points following injection.**

|  | **Subject Name** | **2101 (High dose)** | | | | **1201(Low dose)** | | | **2201(Low dose)** | | |
| --- | --- | --- | --- | --- | --- | --- | --- | --- | --- | --- | --- |
|  | **Time Point** | **Before Injection** | **Day 7** | **Day 28** | **Day 56** | **Before Injection** | **Day 7** | **Day 28** | **Before Injection** | **Day 7** | **Day 28** |
| **Coagulation** | PT (s) | 9.5 | 9.2 | 8.4 | 8.6 | 8.1 | 8.8 | 8.1 | 8.4 | 9 | 8.4 |
|  | APTT (s) | 25.9 | 24.3 | 23 | 23.7 | 24 | 24.9 | 21.9 | 23.4 | 23.1 | 23.6 |
|  | TT (s) | 30.9 | 27.4 | 27.2 | 29.3 | 25.9 | 24.3 | 23.5 | 26.7 | 25.4 | 24.5 |
|  | FIB (g/L) | 1.34 | 3.3 | 1.52 | 1.37 | 2.11 | 3.94 | 2.72 | 2.31 | 4.04 | 2.88 |
| **Serum Biochemical** | ALT (U/L） | 30.9 | 36.3 | 30.2 | 30.3 | 32 | 37.7 | 23.2 | 44.2 | 59.5 | 38 |
|  | AST (U/L） | 42.6 | 55.8 | 55.7 | 34 | 32.7 | 41.4 | 36.8 | 38.5 | 56.2 | 49.8 |
|  | GGT (U/L） | 51.3 | 35.6 | 37.7 | 47.1 | 118.8 | 74.6 | 82 | 100.4 | 76.3 | 81.6 |
|  | ALP (U/L） | 413.9 | 418.1 | 307.4 | 377.9 | 635.7 | 517.1 | 449.3 | 626.4 | 463.8 | 408 |
|  | CK (U/L） | 198.5 | 163.2 | 258.6 | 201.1 | 94.2 | 121.4 | 104.8 | 235 | 182.3 | 226.5 |
|  | TG (mm/L） | 0.79 | 0.37 | 0.53 | 0.41 | 0.33 | 0.43 | 0.22 | 0.22 | 0.29 | 0.24 |
|  | Cr (µm/L） | 59.3 | 54.8 | 51.9 | 60 | 60.7 | 56.6 | 51 | 63.4 | 50.7 | 49.7 |
|  | BUN (mm/L） | 4.9 | 4.8 | 4.3 | 4.1 | 4.9 | 4.6 | 5.2 | 5.4 | 6.1 | 5.6 |
|  | GLUC (mm/L） | 3.37 | 4.68 | 4.03 | 4.48 | 3.59 | 3.44 | 3.31 | 3.72 | 3.28 | 2.57 |
|  | TBil (µm/L） | 3.2 | 1.1 | 0.9 | 1 | 2.7 | 1.5 | 2.1 | 1.1 | 0.3 | 0.9 |
|  | TP (g/L） | 74.7 | 71.6 | 75 | 77.5 | 72.6 | 67.8 | 70.7 | 79.9 | 73 | 78.7 |
|  | ALB (g/L） | 49.2 | 42 | 44.6 | 46.6 | 48.2 | 41.3 | 44.1 | 46.3 | 39.4 | 41.5 |
|  | TC (mm/L） | 3.82 | 3.83 | 3.54 | 4.22 | 3.5 | 3.23 | 3.04 | 2.88 | 2.73 | 2.74 |
|  | C3 (g/L） | 1.06 | 1.6 | 1.07 | 1.1 | 1 | 1.34 | 1.1 | 1.14 | 1.55 | 1.27 |
|  | IgM (g/L） | 1.294 | 1.08 | 1.247 | 1.389 | 0.898 | 0.955 | 0.84 | 1.564 | 1.633 | 1.531 |
|  | IgG (g/L） | 11.7 | 10.36 | 12.41 | 12.21 | 7.36 | 6 | 6.63 | 11.97 | 9.87 | 12.87 |
|  | IgA (g/L） | 1.11 | 1.27 | 1.25 | 1.25 | 2.98 | 2.98 | 3.06 | 3.84 | 3.8 | 4.35 |
|  | C4 (g/L） | 0.2 | 0.45 | 0.22 | 0.22 | 0.17 | 0.36 | 0.29 | 0.17 | 0.33 | 0.32 |
| **Hematology** | WBC (^9/L） |  | 10.28 | 11.54 | 9.34 | 11.37 | 13.99 | 14.58 | 7.87 | 14.07 | 9.79 |
|  | RBC (^12/L） | 5.22 | 4.42 | 4.81 | 5.76 | 6.3 | 5.6 | 5.38 | 4.41 | 3.72 | 4.44 |
|  | Hb (g/L） | 133 | 112 | 127 | 142 | 144 | 124 | 124 | 116 | 99 | 115 |
|  | HCT(%) | 41.2 | 36.5 | 41.9 | 49.1 | 42.8 | 38.6 | 40.3 | 37.3 | 31.6 | 36.9 |
|  | MC (/fL) | 79 | 82.6 | 87.1 | 85.3 | 68 | 68.9 | 74.8 | 84.5 | 84.8 | 83.1 |
|  | MCH (pg) | 25.5 | 25.4 | 26.5 | 24.7 | 22.8 | 22.1 | 23.1 | 26.3 | 26.5 | 26 |
|  | MCHC (g/L） | 323 | 307 | 304 | 289 | 336 | 321 | 309 | 311 | 312 | 313 |
|  | RDW(%) | 14.7 | 15.2 | 15 | 13.9 | 16.8 | 16.8 | 17.5 | 14.7 | 14 | 13.6 |
|  | PLT (^9/L） | 333 | 496 | 405 | 403 | 410 | 559 | 450 | 350 | 406 | 398 |
|  | MPV (fL) | 9 | 8.7 | 9.1 | 8.8 | 9.8 | 9.6 | 9.7 | 9 | 8.6 | 8.8 |
|  | PDW (%) | 43.6 | 49.8 | 48.5 | 44.6 | 74.9 | 83 | 70.7 | 48.1 | 51.7 | 48.4 |
|  | PCT (%) | 0.3 | 0.43 | 0.37 | 0.36 | 0.4 | 0.54 | 0.44 | 0.31 | 0.35 | 0.35 |
|  | NEUT% | 71.9 | 53.1 | 51.6 | 30.3 | 20.8 | 33.1 | 47.3 | 40.3 | 54.4 | 23.6 |
|  | LYMPH% | 25.4 | 42.5 | 43.5 | 63.1 | 74.1 | 60.7 | 44.6 | 54.4 | 39.5 | 68.4 |
|  | MONO% | 1.7 | 2.8 | 2.5 | 4 | 3.7 | 3.9 | 6.3 | 3.6 | 4.6 | 3.5 |
|  | EOS% | 0.5 | 0.9 | 1.7 | 1.8 | 0.4 | 1.2 | 1.2 | 0.5 | 0.5 | 3.2 |
|  | BASO% | 0.2 | 0.2 | 0.3 | 0.4 | 0.7 | 0.3 | 0.3 | 0.4 | 0.2 | 0.6 |
|  | LUC% | 0.3 | 0.5 | 0.4 | 0.5 | 0.4 | 0.7 | 0.4 | 0.8 | 0.8 | 0.7 |
|  | NEUT (^9/L） | 7.38 | 5.46 | 5.95 | 2.83 | 2.36 | 4.63 | 6.89 | 3.17 | 7.66 | 2.31 |
|  | LYMPH (^9/L） | 2.61 | 4.37 | 5.02 | 5.89 | 8.42 | 8.49 | 6.5 | 4.28 | 5.56 | 6.7 |
|  | MONO (^9/L） | 0.17 | 0.29 | 0.29 | 0.37 | 0.42 | 0.55 | 0.91 | 0.28 | 0.64 | 0.34 |
|  | EOS (^9/L） | 0.05 | 0.09 | 0.19 | 0.16 | 0.04 | 0.17 | 0.17 | 0.04 | 0.06 | 0.31 |
|  | BASO (^9/L） | 0.02 | 0.02 | 0.03 | 0.04 | 0.08 | 0.05 | 0.05 | 0.03 | 0.03 | 0.06 |
|  | LUC (^9/L） | 0.03 | 0.05 | 0.05 | 0.05 | 0.04 | 0.1 | 0.05 | 0.06 | 0.11 | 0.07 |
|  | RETIC% | 1.74 | 6.01 | 2.98 | 2.44 | 0.54 | 1.11 | 1.74 | 1.24 | 2.02 | 1.59 |
|  | RETIC (^9/L） | 90.7 | 265.6 | 143.3 | 140.8 | 33.7 | 62.2 | 93.6 | 54.8 | 75.1 | 70.8 |

PT, prothrombin time; APTT, activated partial thromboplastin time; TT, thrombin time; FIB, fibrinogen; ALT, Alanine Aminotransferase; AST, Aspartate Aminotransferase; GGT, Gamma-Glutamyl Transferase; ALP, Alkaline Phosphatase; CK, Creatine Kinase; TG, Triglycerides; Cr, Creatinine; BUN, Urea; GLU, Glucose; TBil, Total Bilirubin; TP, Total Protein; ALB, Albumin; TC, Total Cholesterol; C3, Complement C3; IgM, Immunoglobulin M; IgG, Immunoglobulin G; IgA, Immunoglobulin A; C4, Complement C4; WBC, White Blood Cell Count; RBC, Red Blood Cell Count; Hb, Hemoglobin Concentration; HCT, hematocrit; MCV, Mean Corpuscular Volume; MCH, Mean Corpuscular Hemoglobin; MCHC, Mean Corpuscular Hemoglobin Concentration; RDW, Red Cell Distribution Width; PLT, Platelet Count; MPV, Mean Platelet Volume; PDW, Platelet Distribution Width; PCT, Platelet Crit; NEUT%, Neutrophil Percentage; LYMPH%, Lymphocyte Percentage; MONO%, Monocyte Percentage; EOS%, Eosinophil Percentage; BASO%, Basophil Percentage; LUC%, Large Unstained Cells Percentage; NEUT, Neutrophil Count; LYMPH, Lymphocyte Count; MONO,Monocyte Count; EOS, Eosinophil Count; BASO, Basophil Count; LUC, Large Unstained Cells Count; RETIC%, Reticulocyte Percentage; RETIC, Reticulocyte Count.
